# Sea foams are ephemeral hotspots for distinctive bacterial communities contrasting sea-surface microlayer and underlying surface water

**DOI:** 10.1101/820696

**Authors:** Janina Rahlff, Christian Stolle, Helge-Ansgar Giebel, Nur Ili Hamizah Mustaffa, Oliver Wurl, Daniel P. R. Herlemann

## Abstract

The occurrence of foams at oceans’ surfaces is patchy and generally short-lived but a detailed understanding of bacterial communities inhabiting sea foams is lacking. Here we investigated how marine foams differ from the sea-surface microlayer (SML), a <1 mm thick layer at the air-sea interface and underlying water from 1 m depth. Samples of sea foams, SML and underlying water collected from the North Sea and Timor Sea indicated that foams were often characterized by a high abundance of small eukaryotic phototrophic and prokaryotic cells as well as a high concentration of surface-active substances (SAS). Amplicon sequencing of 16S rRNA (gene) revealed a distinctive foam bacterial community compared to SML and underlying water, with high abundance of *Gammaproteobacteria*. Especially *Pseudoalteromonas* and *Vibrio*, typical SML dwellers, were highly abundant, active foam inhabitants and thus might enhance foam formation and stability by producing SAS. Despite a clear difference in the overall bacterial community composition between foam and SML, the presence of SML bacteria in foams supports previous assumptions that foam is strongly influenced by the SML. We conclude that active and abundant bacteria from interfacial habitats potentially contribute to foam formation and stability, carbon cycling and air-sea exchange processes in the ocean.

**One-sentence summary:** Floating foams at the oceans’ surface have a unique bacterial community signature in contrast to sea-surface microlayer and underlying water but receive and select for bacterial inhabitants from surface habitats.

## Introduction

Foams are patches floating on the water surface and may appear in any aquatic habitat. Foam is loosely defined as a dispersion of gas in liquid in the presence of surface-active substances (SAS) (Schilling and Zessner 2011). Convergence at zones of downwelling water and fronts, currents, and breaking waves compress SAS and lead to foam formation at the sea surface and occasionally cause massive foam aggregates at beaches and in coastal zones (Bärlocher, *et al*. 1988; Eisenreich, *et al*. 1978; Jenkinson, *et al*. 2018; Kesaulya, *et al*. 2008; Thornton 1999). Furthermore, rising bubbles that do not burst immediately but accumulate at the surface can cause foam formation (Schilling and Zessner 2011). The nature, distribution and occurrence of foam in the marine environment is elusive, since its lifespan is limited to hours or days (Pugh 1996; Velimirov 1980), and the mean coverage of the ocean’s surface by foams (white caps) is 1 - 6% based on satellite observations (Anguelova and Webster 2006).

One major prerequisite for foam formation are SAS, representing a complex mixture of mainly organic compounds. Due to their amphipathic nature, SAS accumulate at the sea surface (Wurl, *et al*. 2009) and influence CO2 air-sea gas exchange (Pereira, *et al*. 2018; Ribas-Ribas, *et al*. 2018). In foams, SAS can originate from a variety of sources such as marine bacteria (Satpute, *et al*. 2010), kelp mucilage (Velimirov 1980), exudates of alive or broken phytoplankton cells (Frew, *et al*. 1990; Velimirov 1980; Velimirov 1982; Wegner and Hamburger 2002), or other organic detritus (Velimirov 1980). In addition, organic materials such as biogenic lipids and amino acids that accumulate at the sea surface during phytoplankton blooms, are important substrates for the formation of foam (Eberlein, *et al*. 1985; Hunter, *et al*. 2008; Riebesell 1993). Even if foam is generally short-lived, its high concentration of organic matter (Eisenreich, *et al*. 1978; Johnson, *et al*. 1989), especially of proteins and carbohydrates (Stefani, *et al*. 2016), allows these nutrient-rich islands to function as microbial habitats. Despite being ephemeral feeding grounds, foams are remarkably rich and diverse in microorganisms (Tsyban 1971), including bacteria (Gobalakrishnan, *et al*. 2014), protists, and algae (Harold and Schlichting 1971; Maynard 1968). In addition, foams were shown to enclose copepods, polychaete, and tunicate larvae (Armonies 1989; Castilla, *et al*. 2007) and to trap microalgae (Roveillo, *et al*. 2020), thus forming potentially important food sources for the higher trophic levels of the food web (Bärlocher, *et al*. 1988; Craig, *et al*. 1989; Scully 2009). In addition, a pharmaceutical potential of sea foam inhabiting microbes was recently suggested (Oppong-Danquah, *et al*. 2020).

The sea-surface microlayer (SML) is a <1 mm thick, biofilm-like layer (Wurl and Holmes 2008; Wurl, *et al*. 2016), located at the air-sea boundary of all aquatic ecosystems (Supplementary Figure 1). It is characterized by remarkably different physicochemical and biological properties that allow its differentiation from the underlying water (ULW) (Cunliffe, *et al*. 2013; Hardy 1982). Research throughout the last decades revealed that the accumulation of inorganic and organic substances and particles (including microorganisms) at the sea surface is a widespread phenomenon with important implications for carbon cycling processes (Engel, *et al*. 2017; Rahlff 2019; Wurl, *et al*. 2017). The interfacial position of the SML represents a challenging environment for inhabiting organisms also known as neuston (Naumann 1917). Differences in bacterial community composition between SML and ULW have been related to meteorological conditions (Agogué, *et al*. 2005b; Rahlff, *et al*. 2017a; Stolle, *et al*. 2011), however the specific adaptation of bacteria to the SML habitat remains an open question (Agogué, *et al*. 2005a).

**Figure 1:**
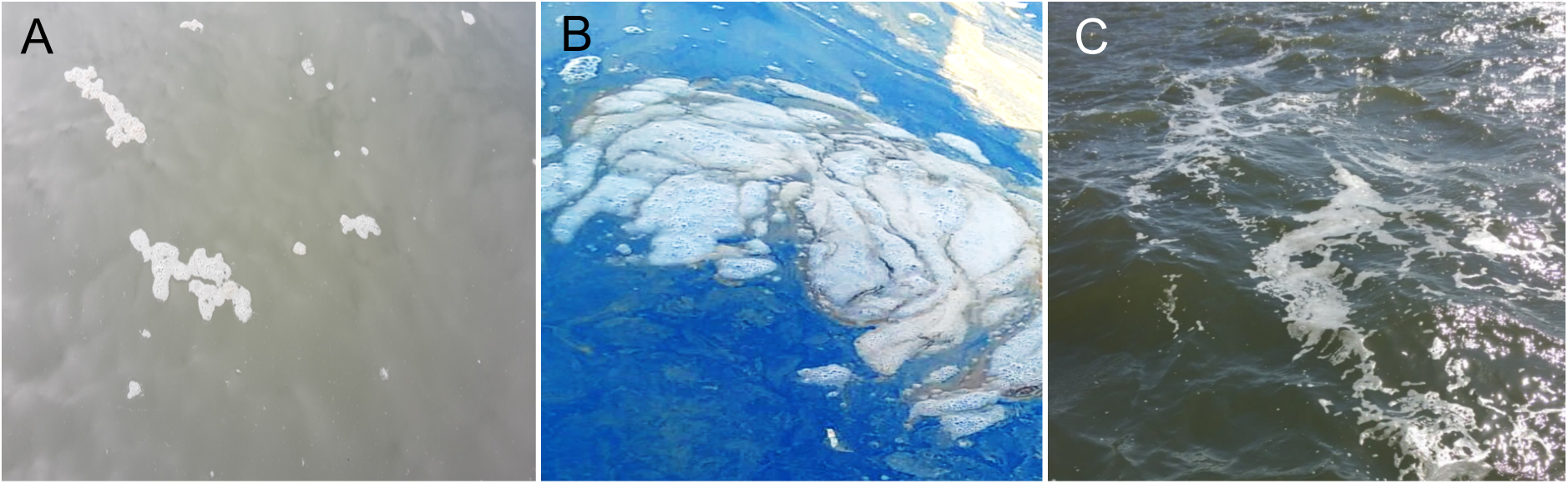
Marine foam originating from A) presumptive phytoplankton exudates (Jade Bay), B) *Trichodesmium* bloom (Timor Sea) and C) whitecaps produced by convergence of surface water (Jade Bay, North Sea).

Neuston and plankton have a demonstrated role in gas exchange across air-water interfaces (Calleja, *et al*. 2005; Upstill-Goddard, *et al*. 2003). Moreover, the impact of the neuston on carbon cycling can be high due to its higher abundance of bacteria compared to the ULW (Hardy 1982). This includes enhanced heterotrophic activity (Obernosterer, *et al*. 2005), turnover of organic matter (Reinthaler, *et al*. 2008), colonization of buoyant and sinking particle aggregates (Bigg, *et al*. 2004; Leck and Bigg 2017), as well as viral lysis of prokaryotic cells (Ram, *et al*. 2018). First approaches to quantify the contributions of neuston activity to underlying water oxygen concentrations revealed maximum neuston contributions of ≤7.1% (Rahlff, *et al*. 2019).

Napolitano and Cicerone (1999) suggested that 1 L of foam water represents 2 m^2^ of SML of 50 µm thickness, i.e. foams are essentially compressed SML. Supporting this idea, enrichment of bacteria in foams compared to SML and/or ULW has been reported (Kuznetsova and Lee 2002; Rahlff, *et al*. 2017a; Tsyban 1971). In addition, during blooms of *Trichodesmium erythraeum*, high concentrations of this species were also found in the corresponding foam (Maynard 1968). However, a thorough characterization of foam microbial community composition compared to the SML and ULW has never been performed. Using a microscopic approach, Druzhkov, *et al*. (1997) found a highly identical taxonomic composition of heterotrophs (nanoflagellates <5 µm and bacteria), nano- and microphytoplankton in foam and the SML. The authors further described higher abundances (one order of magnitude) of autotrophs, but not of heterotrophs in foams compared to the SML.

In this study, we investigated the bacterial community composition of marine foams in direct comparison to SML and ULW. Based on the theory that foam is an extreme condensed form of the SML (Napolitano and Cicerone 1999), we hypothesized that the bacterial community composition of foam and SML are more similar than between foam and ULW. Since the SML is also considered an extreme habitat (Maki 1993) likely comprising many dead or dormant cells, we differentiated between active and total bacteria as inferred from a complementary DNA (cDNA) and DNA-based 16S rRNA amplicon sequencing, respectively. Overall, we provide a detailed understanding of the bacterial community composition associated with marine foams in comparison to SML and ULW with likely implications for foam formation and stability, air-sea exchange processes and carbon cycling.

## Materials & Methods

### Field sample collection

Field sampling was conducted from the bow of a small boat in the Jade Bay, North Sea offshore Wilhelmshaven, Germany (Supplementary Table 1) in spring and summer 2016. Foams originated from different sources such as phytoplankton exudates and convergence of surface water (Figure 1A, Figure 1C, Supplementary Table 1). Additional samples were collected during a *Trichodesmium* sp. bloom encountered in the Timor Sea at stations 4, 5b and 8 (Figure 1B, Supplementary Table 1) in October 2016 during *R/V* Falkor cruise FK161010 as described by Wurl, *et al*. (2018). Foam, and corresponding SML and ULW samples (here referred to as ‘set’) were collected from each location. From the Jade Bay, six sets were sampled in total, i.e., two from each month in April, May and July 2016. From the Timor Sea, one set was collected from each of three stations, but sequencing was only done for station 8, i.e., one sampling set. Foam and SML were collected using the glass plate technique (Harvey and Burzell 1972) with a withdrawal rate of 5-6 cm s^-1^ as suggested by Carlson (1982). The glass plate was cleaned with 70% ethanol and rinsed with sample water before use. Material adhering on the glass plate was removed by wiping its surface with a squeegee into a sample-rinsed brown bottle. The procedure was repeated until the required volume of approximately 100 mL was collected (∼20 dips). SML samples were collected between the foam patches and any dips contaminated with foam were rejected, and the glass plate was cleaned with ethanol again. Collected foams were not generated by the small boat whose engine was turned off. Samples from the ULW were taken at a depth of 1 m around the foams by using a syringe connected to a hose. All samples were kept on ice and immediately processed after sampling, since Velimirov (1980) showed that bacterial density in old foam was significantly higher than in fresh foam.

**Table 1:** Absolute and relative abundances given as enrichment factor (EF) of prokaryotes, small phototrophic eukaryotes and surface-active substances (SAS) in foam (F), sea-surface microlayer, SML (S) and underlying water, ULW (U), NS=North Sea, NA=not available, Teq=Triton X-100 equivalents, TS=Timor Sea

|  | Foam | SML | ULW | EF (F/S) | EF (F/U) | EF (S/U) |
| --- | --- | --- | --- | --- | --- | --- |
| **Prokaryotes (cells mL<sup>-1</sup>)** | **Absolute values (10<sup>6</sup> cells mL<sup>-1</sup>)** |  |  | **Relative values** |  |  |
| NS_St1_210416 | 4.89 | NA | 2.56 | NA | 1.9 | NA |
| NS_St2_210416 | 2.63 | 2.62 | 2.48 | 1.0 | 1.1 | 1.1 |
| NS_St1_190516 | 13.70 | 3.34 | 3.23 | 4.1 | 4.2 | 1.0 |
| NS_St3_190516 | 46.20 | 4.57 | 3.13 | 10.1 | 14.8 | 1.5 |
| NS_St1_190716 | 6.61 | 3.90 | 3.71 | 1.7 | 1.8 | 1.1 |
| NS_St2_190716 | 4.99 | 3.39 | 3.48 | 1.5 | 1.4 | 1.0 |
| TS_St4_151016 | 9.97 | 1.77 | 1.07 | 5.6 | 9.3 | 1.7 |
| TS_St5b_171016 | 33.90 | NA | 1.01 | NA | 33.6 | NA |
| TS_St8_191016 | 5.83 | 0.98 | 1.18 | 5.9 | 4.9 | 0.8 |
| **Small phototrophic eukaryotes (cells mL<sup>-1</sup>)** | **Absolute values (10<sup>4</sup> cells mL<sup>-1</sup>)** |  |  | **Relative values** |  |  |
| NS_St1_210416 | 1.85 | 1.03 | 1.61 | 1.8 | 1.1 | 0.6 |
| NS_St2_210416 | 1.38 | 1.41 | 2.24 | 1.0 | 0.6 | 0.6 |
| NS_St1_190516 | 29.60 | 0.85 | 2.30 | 34.8 | 12.9 | 0.4 |
| NS_St3_190516 | 57.10 | 1.02 | 1.88 | 56.0 | 30.4 | 0.5 |
| NS_St1_190716 | 9.10 | 3.97 | 4.17 | 2.3 | 2.2 | 1.0 |
| NS_St2_190716 | 4.23 | 2.49 | 2.82 | 1.7 | 1.5 | 0.9 |
| TS_St4_151016 | 2.14 | 0.37 | 0.11 | 5.8 | 20.2 | 3.5 |
| TS_St5b_171016 | 10.50 | NA | 0.13 | NA | 81.4 | NA |
| TS_St8_191016 | 2.73 | 0.12 | 0.26 | 23.7 | 10.4 | 0.4 |

**SAS (µg Teq L<sup>-1</sup>)**
|  | **Absolute values** |  |  | **Relative values** |  |  |
| --- | --- | --- | --- | --- | --- | --- |
| NS_St1_210416 | NA | NA | NA | NA | NA | NA |
| NS_St2_210416 | NA | NA | NA | NA | NA | NA |
| NS_St1_190516 | 77496 | 576 | 213 | 134.4 | 364.5 | 2.7 |
| NS_St3_190516 | 148233 | 1753 | 223 | 84.6 | 665.0 | 7.9 |
| NS_St1_190716 | 900 | 716 | 180 | 1.3 | 5.0 | 4.0 |
| NS_St2_190716 | 1397 | 270 | 133 | 5.2 | 10.5 | 2.0 |
| TS_St4_151016 | 69370 | 240 | 268 | 288.9 | 258.5 | 0.9 |
| TS_St5b_171016 | 67546 | 66 | 109 | 1020.5 | 618.7 | 0.6 |
| TS_St8_191016 | 28797 | 255 | 171 | 113.1 | 168.5 | 1.5 |

### Concentration of surface-active substances

The concentration of surface-active substances (SAS) was measured by the voltammetry VA Stand 747 (Methrom, Herisau, Switzerland) with a hanging drop mercury electrode as previously described (Ćosović and Vojvodić 1998; Wurl, *et al*. 2011). The quantification is based on SAS adsorption on the Hg electrode measured by the change of capacity current (ΔIc) at an applied potential (E) of -0.6 V (Ćosović and Vojvodić 1998). Before measurement, concentrated samples such as foam samples were diluted with artificial seawater (0.55 M of NaCl solution) to achieve measurements within the linear calibration range. A standard addition technique was used with non-ionic surfactant Triton X-100 (Sigma Aldrich, Taufkirchen, Germany) as a standard. SAS concentration in the samples was measured using two to three analytical replicates, resulting in relative standard deviations below 6% (Rickard, *et al*. 2019). Concentration of SAS is expressed as the equivalent concentration of the additional Triton X-100 (µg Teq L^−1^).

### Determination of microbial abundance

For determination of prokaryotic and small (< 50 µm) eukaryotic phototrophic cell numbers in all Jade Bay samples and for three stations from the Timor Sea, foam and water samples were fixed with glutaraldehyde (1% final concentration), incubated at room temperature for 1 hour, and stored at -80°C until further analysis. Prior staining and counting by flow cytometry, the particle-enriched foam samples were pre-filtered by gravity onto CellTrics® 50 µm filter (Sysmex Partec, Münster, Germany) to avoid clogging of the instrument by particulate matter. Autofluorescence analysis was used to count small eukaryotic phototrophic cells (Marie, *et al*. 2000), and prokaryotic cells were stained with SYBR® Green I Nucleic Acid Gel Stain (9x in final concentration, Thermo Fisher Scientific, Darmstadt, Germany) following a protocol of Giebel, *et al*. (2019). We did not measure biological replicates of samples because the coefficient of variation for SML and ULW flow cytometry samples is <5% (Rahlff, *et al*. 2017a).

### Calculation of Enrichment Factors (EF)

Enrichment factors (EF) of SAS and cell counts were calculated for the pairings foam/SML (F/S), foam/ULW (F/U) and SML/ULW (S/U) (Table 1). Since the concentration of SAS or the abundance of cells in a foam or SML sample was divided by its SML or ULW counterpart, an EF>1 implies enrichment of the parameter, whereas an EF<1 indicates depletion.

### Nucleic acid extraction and PCR

A two-step filtration of foam, SML and ULW samples was conducted as performed in previous studies (Garneau, *et al*. 2009; Stolle, *et al*. 2011). Sample water was filtered through 3 µm pore size (particle-associated cells) polycarbonate filters, after which the filtrate was filtered onto 0.2 µm pore size (free-living cells) polycarbonate filters (Merck Millipore, Darmstadt, Germany). Foam from the Timor Sea (Station 8) collected during a bloom of *Trichodesmium* sp. was additionally pre-filtered on a 100 µm mesh before subsequent filtration on the 3 µm pore size filter. All filters were initially stored at -80°C prior analysis. Extraction of DNA and RNA from the filters was performed by using the DNA + RNA + Protein Extraction Kit (Roboklon, Berlin, Germany) with a modified protocol (Rahlff, *et al*. 2017a). Remaining DNA in RNA samples was digested on-column using 3 U of DNase and subsequently checked for contaminations with genomic DNA by PCR (Supplementary Figure 2). A quantity of 10 ng RNA was reversely transcribed to cDNA using the NG dART Kit (Roboklon, Berlin, Germany) including negative controls either without reverse transcriptase or without RNA (Supplementary Figure 2). The reaction was incubated using the primer 1492R (5’-GGTTACCTTGTTACGACTT-3’, adapted from (Lane 1991)) for 60 minutes at 50°C followed by 5 minutes at 85°C. All DNAs and cDNAs were quantified using the Quant-iT™ PicoGreen™ dsDNA assay (Thermo Fisher Scientific, Darmstadt, Germany).

**Figure 2:**
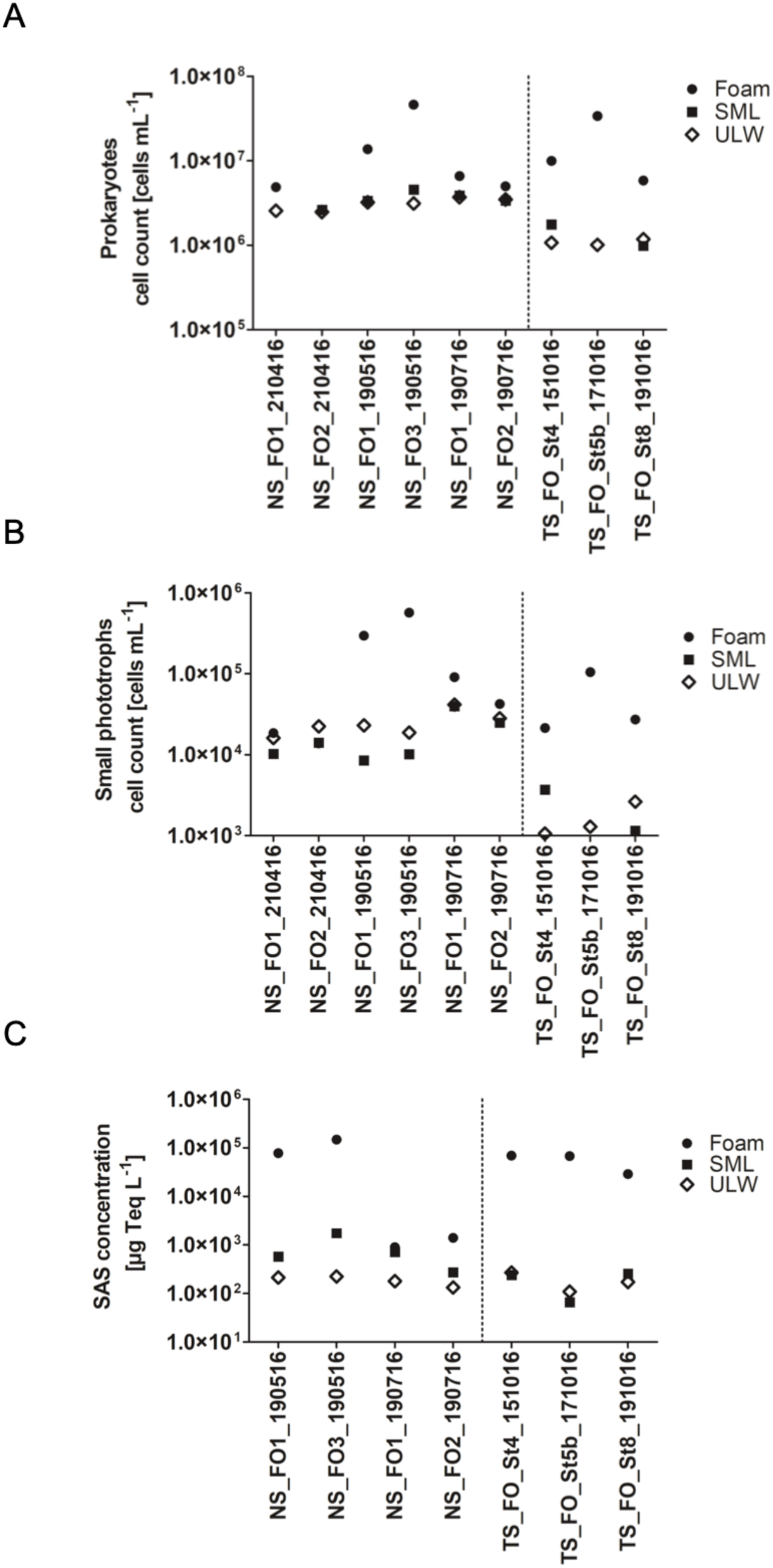
Absolute cell counts mL^-1^ for A) prokaryotes and B) small eukaryotic phototrophic cells and C) concentration of surface-active substances (SAS) in µg Teq L^-1^ for foam, sea-surface microlayer (SML) and underlying water (ULW). Foam samples from the North Sea NS_FO1_210416, and NS_FO2_210416 were produced by waves, whereas NS_FO1_190516, NS_FO2_190516 and NS_FO1_190716 are foams likely derived from phytoplankton exudates. Foams from the Timor Sea (TS) originated from a *Trichodesmium* sp. bloom.

### 16S rRNA library preparation, sequencing run and data analysis

The bacterial 16S rRNA gene was amplified using Bakt_341F (5’-CCTACGGGNGGCWGCAG-3’) and Bakt_805R (5’-GACTACHVGGGTATCTAATCC-3’) (Herlemann, *et al*. 2011) with the following modifications. Genomic DNA was amplified with 35 cycles prior Index-PCR. The cDNA samples were amplified with 25 cycles prior Index-PCR. Amplicon PCR, Index PCR, quantity and quality control and sequencing of the individual libraries as pool in one Illumina MiSeq run was performed by an external provider (Eurofins Genomics, Ebersberg, Germany). Raw sequencing data were deposited at the European Nucleotide Archive (ENA) under accession number PRJEB34343. For data analysis, the resulting sequences were assembled using QIIME 1.9.1 (Caporaso, *et al*. 2010) “joins paired-end Illumina reads” function with default settings to merge forward and reverse sequence with an overlap of at least 30 bp. Sequences without overlap were removed. After converting fastq to fasta using the “convert_fastaqual_fastq” function the resulting sequences were evaluated using the SILVA NGS pipeline. The SILVA next - generation sequencing (NGS) pipeline (Glöckner, *et al*. 2017) performs additional quality checks according to the SINA-based alignments (Pruesse, *et al*. 2012) with a curated seed database in which PCR artifacts or non-SSU reads are excluded (based on SILVA release version 128 (Pruesse, *et al*. 2007). The longest read serves as a reference for the taxonomic classification in a BLAST (version 2.2.28+) search against the SILVA SSU Ref dataset. The classification of the reference sequence of a cluster (98% sequence identity) is mapped to all members of the respective cluster and to their replicates. Best BLAST hits were only accepted if they had a (sequence identity + alignment coverage)/2 ≥ 93% or otherwise defined as unclassified. SILVA NGS classified a total of 9182084 reads (2% were rejected by the quality control). Sequences assigned to chloroplasts, mitochondria, eukaryotes and *Archaea* were removed since the primer set employed in the analysis has only a very limited coverage of these groups.

### Statistical analyses

Operational taxonomic unit (OTU) counts based on genus level were rarefied to 43,500 reads per sample using the single_rarefraction.py script implemented in QIIME. Venn diagrams were calculated using the ugent webtool (http://bioinformatics.psb.ugent.be/webtools/Venn/). We visualized the differences in the bacterial community composition through non-metric multidimensional scaling (NMDS) plots using Bray–Curtis dissimilarity indices based on a genus rank classification. A linear discriminant analysis effect size (LEfSe) analysis was performed to determine bacterial groups which are significantly different between the samples using the ‘one against all’ strategy for multi-class analysis (Segata, *et al*. 2011). The program LEfSe uses a non-parametric test that combines standard tests for statistical significance with additional tests encoding biological consistency and effect relevance. *P*<0.05 was regarded as statistical significance.

Differences in the number of 16S rRNA and 16S rRNA gene-derived OTUs between habitats (foam, SML, ULW) and attachment status were statistically analyzed using a Kruskal-Wallis test, and *post hoc* multiple pairwise comparisons were conducted based on Dunn’s z statistic approximations to the actual rank statistics within the package “dunn.test” (Dinno and Dinno 2017) in R version 4.0.3 (Team 2017). The null hypothesis was rejected if *p*>0.05. For the same samples, comparisons on phylum and OTU-based differences were statistically investigated using a one-way Analysis of Similarities (ANOSIM) based on Bray Curtis dissimilarity and 9999 permutations, Bonferroni correction and a significance level of 95%.

## Results

### Foams are enriched with surface-active substances and microorganisms

Overall, foams from both sampling areas (North Sea and Timor Sea), were enriched with prokaryotic microorganisms, small eukaryotic phototrophic cells, and SAS (Table 1). Cell counts of prokaryotic microorganisms ranged between 2.63 - 46.2 x 10^6^, 0.98 – 4.57 x 10^6^, and 1.01 - 3.71 x 10^6^ cells mL^-1^ in foam, SML and ULW, respectively (Figure 2A). Thus, prokaryotic microorganisms in foams were enriched with a maximum EF (enrichment factor) of 10.1 and 5.9 over SML, and with a maximum EF of 14.8 and 33.6 over ULW in North Sea and Timor Sea, respectively (Table 1). However, these numbers likely represent an underestimation of cell counts because pre-filtration of foam samples onto a 50 µm sieve before the flow cytometry measurement likely excluded some bigger aggregates of cells and colloid material. Prokaryotic cells in the SML were enriched with a maximum EF of 1.5 and 1.7 over ULW in North Sea and Timor Sea, respectively. Likewise, the total number of small eukaryotic phototrophic cells, was always higher in foam (range=1.38 - 57.1 x 10^4^ cells mL^- 1^) compared to SML (range=1.15 - 39.7 x 10^3^ cells mL^- 1^) and ULW (range=1.06 - 41.7 x 10^3^ cells mL^- 1^, Figure 2B). Thus, the maximum EF was 3.5 and 81.2 for SML over ULW and foam over ULW, respectively. The absolute number of small eukaryotic phototrophic cells was two orders of magnitude lower compared to the prokaryotic cell counts (Figure 2 A&B). Interestingly, small eukaryotic phototrophic cells were often depleted in the SML compared to the ULW (S/U minimum EF=0.4), while they were enriched in foams over ULW at the same time (F/U EF=12.9 (Table 1)).

Foams contained the highest SAS concentrations compared to the other two habitats (Figure 2C). SAS concentrations in foams varied between 900 to 148,233 µg Teq L^-1^ in North Sea and Timor Sea, whereas SML SAS concentrations were in a range of 66 to 1,753 µg Teq L^-1^, and ULW SAS concentrations in a range of 109 to 223 µg Teq L^-1^ (Table 1). While SAS concentrations in the SML were enriched with a maximum EF of 7.9 compared to ULW, their concentration in foams compared to ULW was typically enriched by two orders of magnitude (EF ranging from 5 to 665).

### Changes in the number of OTUs among foam, sea-surface microlayer and underlying water

We analyzed the bacterial community composition of all North Sea samples and could not detect any obvious difference between sampling dates (Supplementary Figure 3). In the following, we did not differentiate between sampling dates, but by habitat (foam, SML, ULW) attachment status (particle-associated, free-living), and nucleic acid type for 16S rRNA analysis (16S rRNA gene, 16S rRNA) (Figure 3A). Analyses revealed overall higher alpha diversity (numbers of OTUs) in active communities (median=786.5) compared to total communities (median=571).

**Figure 3:**
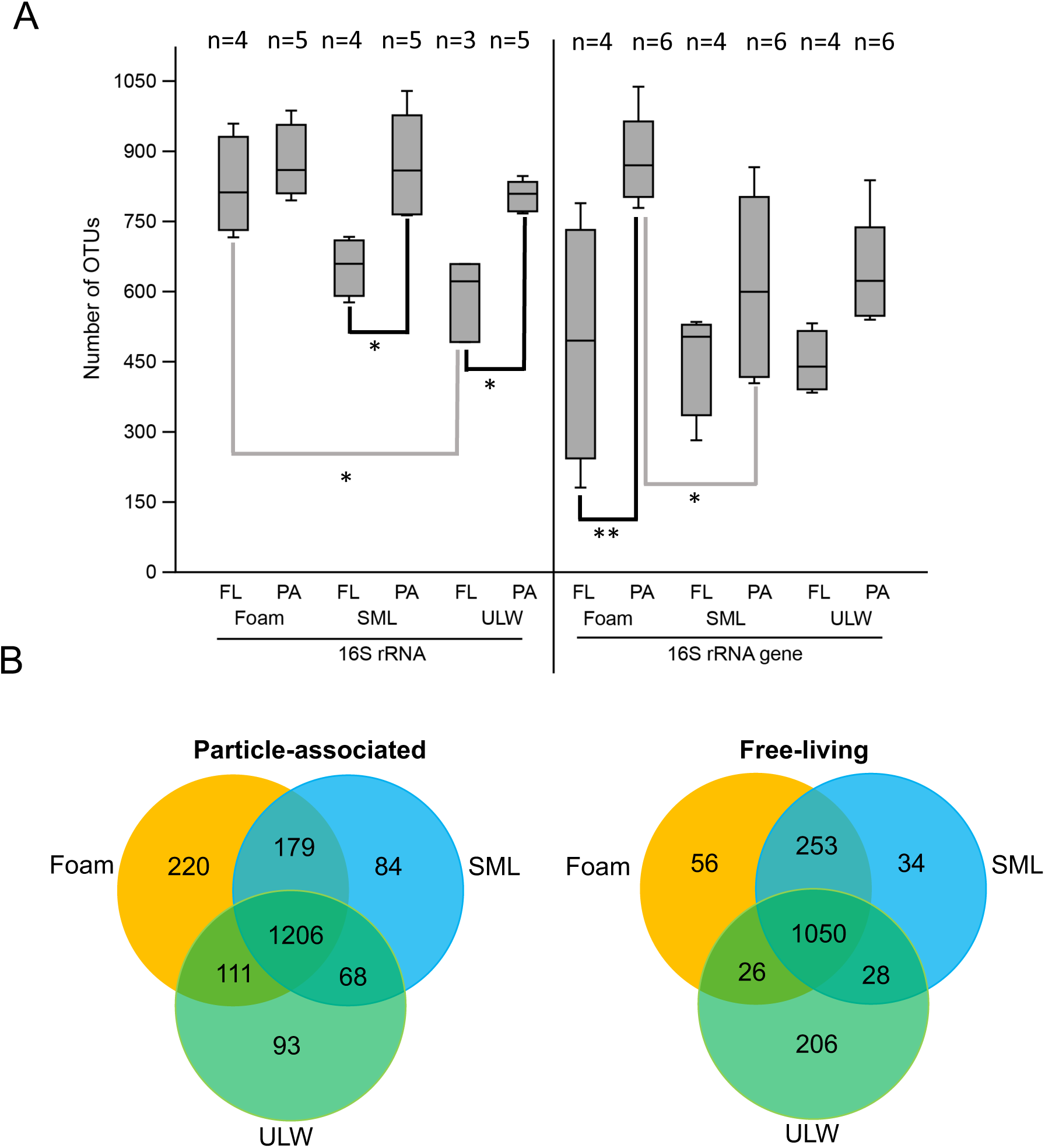
A) 16S rRNA and 16S rRNA gene-derived numbers of operational taxonomic units (OTUs) for foam, sea-surface microlayer (SML) and underlying water (ULW) habitat of pooled North Sea stations. The total number of OTUs of the three habitats is further distinguished between free-living (FL) and particle-associated (PA) bacterial communities. Grey and black lines indicate inter- and intra-habitat comparisons, respectively. The boxplot shows the 25–75 percent quartiles; the median is indicated by the horizontal line inside the box. Error bars show minimal and maximal values. Asterisks indicate the level of significant differences: * p≤0.05, ** p≤0.01; For reasons of the different number of observations (n) see Supplementary Table 1. B) 3-Venn diagram showing overlapping and unique OTUs for foam, SML and ULW separated by free-living and particle-associated OTUs.

In 16S rRNA gene-based samples, the mean number of foam OTUs was significantly increased for particle-associated over free-living communities (Dunn’s test, *p*=0.0090), and also significantly higher compared to the SML particle-associated fraction with (*p*=0.035, Figure 3A). OTU numbers derived from 16S rRNA were significantly different between the particle-associated and the free-living fractions of SML (Dunn’s test, *p*=0.015) and ULW (*p*=0.040) as well as between the free-living communities of foam and ULW (*p*=0.032, Figure 3A). Further significant differences existing between different biomes and different attachment states are shown in Supplementary Table 2. The Shannon-Wiener index (Supplementary Figure 4), which accounts for both abundance and evenness of OTUs, confirmed the above-mentioned trends reflected by the total number of OTUs.

**Figure 4:**
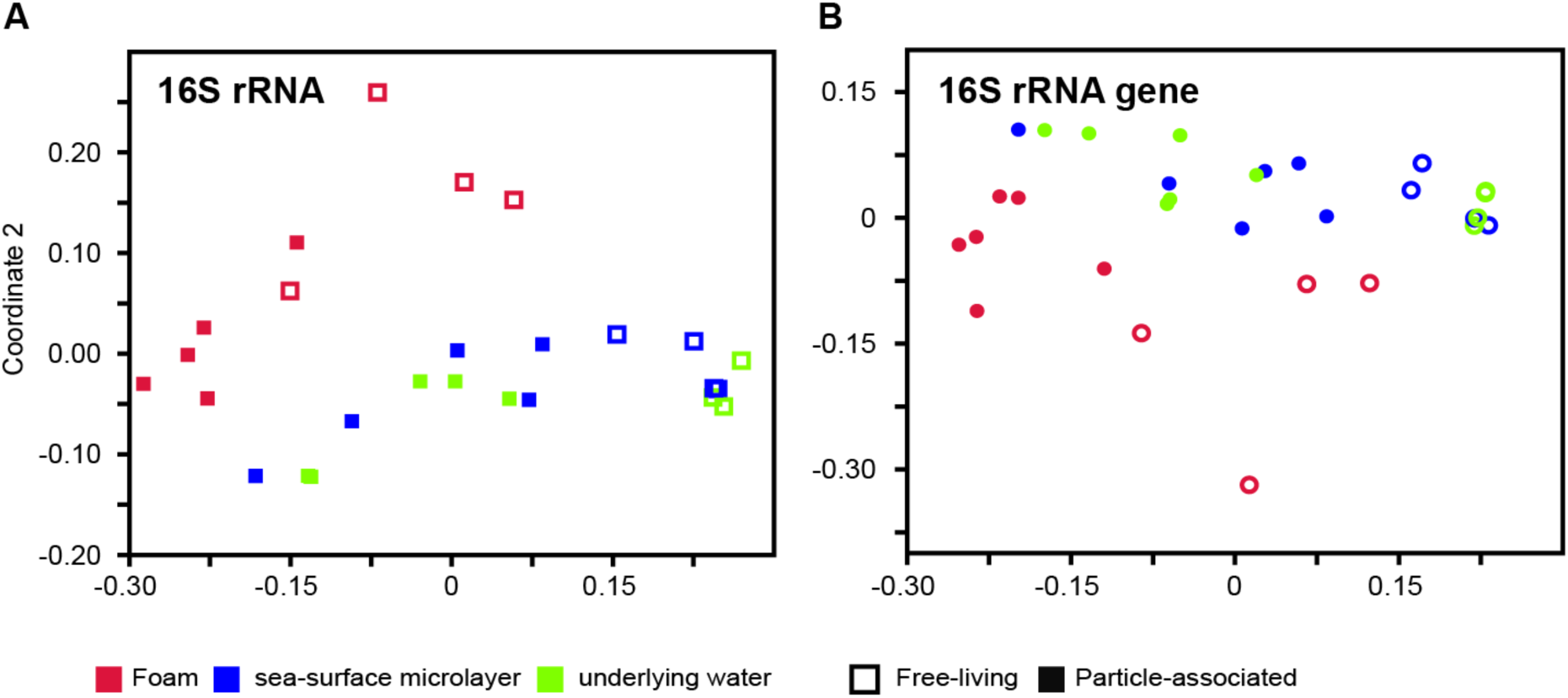
Non-metric multidimensional scaling plot shows distinct clustering of foam (red), sea-surface microlayer (blue) and underlying water (green) bacterial communities. Further separation of communities into A) 16S rRNA-based with free-living (open symbols) and particle-associated (filled symbols) stress=0.14; B) 16S rRNA gene-based with free-living (open symbols) and particle-associated (filled symbols) stress=0.11;

When comparing the 16S rRNA gene-based and 16S rRNA-based as well as the particle-attached and free-living communities, foam, SML and ULW generally shared a high number of similar OTUs (953-1206). Interestingly, foams and SML always had more OTUs in common compared to foam-ULW and ULW-SML (Figure 3B, Supplementary Figure 5). In the particle-attached fraction, foam had the highest fraction of specific OTUs (220) and also shared many with the SML (179). In addition, SML and foam shared many OTUs in the free-living fraction (253) with less specific OTUs in foam (56) and the SML (34) compared to the particle-associated fraction (Figure 3B).

**Figure 5:**
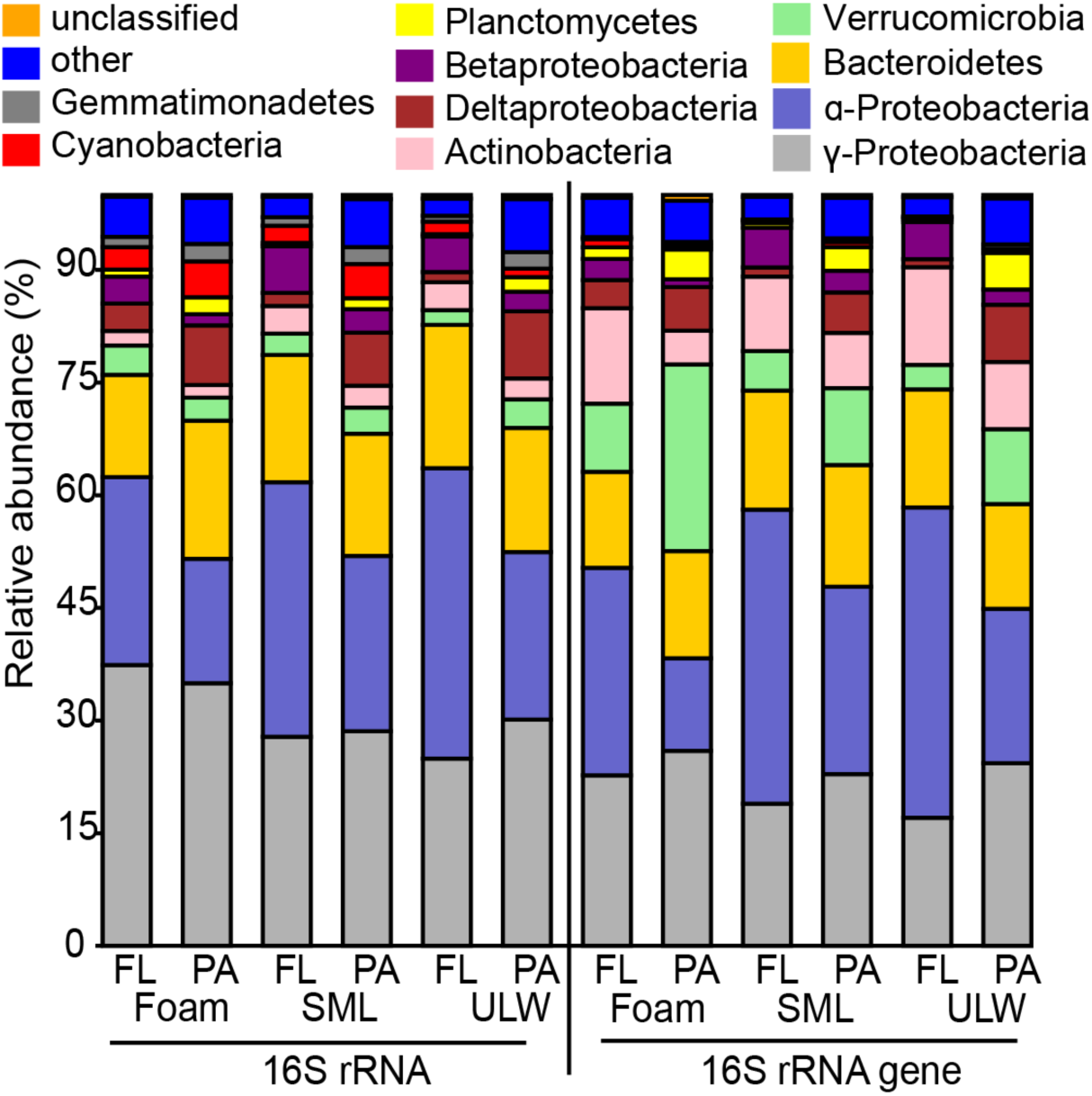
Composition of phylum-level of foam, sea-surface microlayer (SML) and underlying water (ULW) samples of 16S rRNA and 16S rRNA gene-based relative abundance of operational taxonomic units (OTUs) of pooled North Sea stations. Each habitat is further separated into free-living (FL) and particle-associated (PA) bacterial communities.

### Bacterial community composition of sea foam from the Jade Bay, North Sea

Non-metric multidimensional scaling plots comparing the bacterial community composition based on the abundance of OTUs, revealed that the foam bacterial community composition was distinct from SML and ULW communities, irrespective of differentiating 16S rRNA and its gene, or free-living and particle-associated bacterial communities (Figure 4, Supplementary Figure 6). This was supported by the pairwise ANOSIM test, which revealed that the 16S rRNA and 16S rRNA gene-based community differed significantly between foam and SML (both *p*=0.01) as well as between foam and ULW (*p*=0.01 and 0.04, respectively). In contrast, SML and ULW bacterial community composition were more similar to each other as shown by the clustering (Figure 4, Supplementary Figure 6).

**Figure 6:**
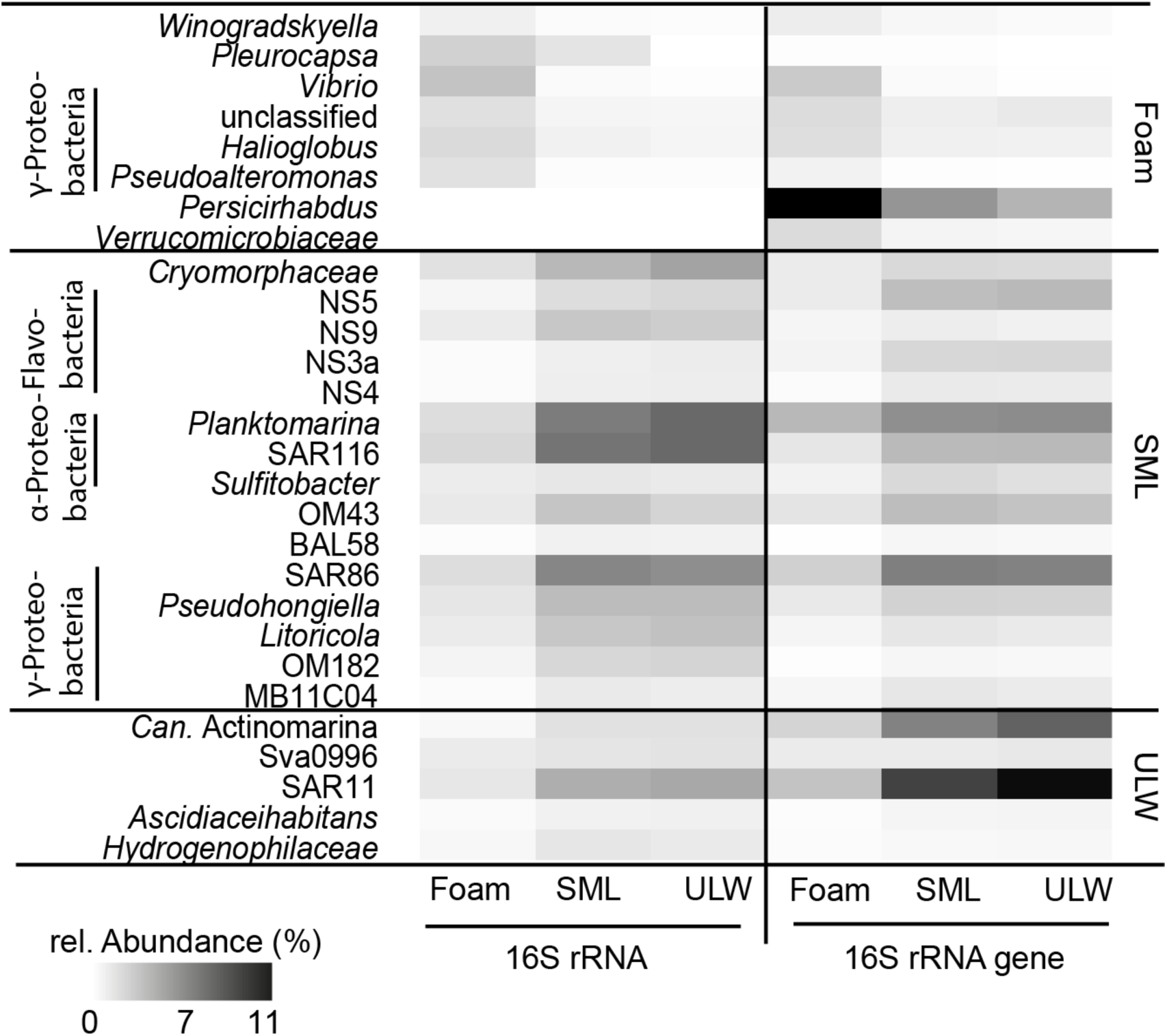
Heat-map showing operational taxonomic units (OTUs) with significant different relative abundance in foam, sea-surface microlayer (SML) and underlying water (ULW) based on LEfSe analysis from pooled North Sea samples; OTUs are derived from sequencing amplicons derived from 16S rRNA and its gene. The vertical axis indicates the key species for these three biomes, respectively, while the horizontal axis shows how abundant these key OTUs were in the other two habitats.

On a phylum-level, 16S rRNA gene-based community analyses showed that for all three habitats *Gammaproteobacteria, Verrucomicrobia* and *Cyanobacteria* formed a higher portion of particle-associated than free-living communities with the exception of *Cyanobacteria* in foam (Figure 5). In contrast, *Alphaproteobacteria* and *Actinobacteria* were more abundant in the free-living fraction (Figure 5, Supplementary Table 3).

Most phyla were similarly distributed in 16S rRNA and 16S rRNA gene-derived bacterial communities, suggesting that many of the total phyla were also active. *Gammaproteobacteria*, as one exception, tended to have higher relative abundance in the 16S rRNA-based community composition in foam (37.4% and 35.0% of free-living and particle-associated OTUs, respectively) compared to the 16S rRNA gene-based community composition (22.7% and 26.0% of free-living and particle-associated OTUs, respectively). In contrast, *Verrucomicrobia* had a higher portion in all habitats and fractions based on 16S rRNA gene compared to 16S rRNA. This was particularly striking for particle-associated communities in foams, where *Verrucomicrobia* were quite abundant among the total community (24.9%) but little active (3.1%). The 16S rRNA-based communities of foam contained less *Alphaproteobacteria* but more *Gammaproteobacteria* compared to SML and ULW communities (Figure 5).

MB11C04 marine group (*Verrucomicrobia*), SAR11 clade (*Alphaproteobacteria*) and *Oceanospirillales* (*Gammaproteobacteria*) had a higher relative abundance in ULW and SML compared to foam based on 16S rRNA gene and 16S rRNA and when comparing the respective attachment status, i.e., free-living and particle-associated fractions. (Supplementary Figures 7, 8, 9). A higher relative abundance of active OTUs (based on 16S rRNA) in foam compared to SML and ULW was found among the *Puniceicoccales* (*Verrucomicrobia*), *Sphingomonadales* (*Alphaproteobacteria*), *Alteromonadales* and *Vibrionales* (both *Gammaproteobacteria*) (Supplementary Figure 7, 8, 9). Active, free-living OTUs belonging to the order *Flavobacteriales* and *Oceanospirillales* were more, whereas free-living *Sphingobacteriales* were less numerous than their particle-associated counterparts in all three habitats (Supplementary Figures 7 & 10).

Except from the order *Rhodobacterales* (Supplementary Figure 9), foam generally had less alphaproteobacterial 16S rRNA gene-based OTUs compared to SML and ULW. However, foam contained a higher 16S rRNA gene-based relative abundance of *Verrucomicrobia* and *Gammaproteobacteria* compared to SML and ULW (Figure 5). Among the *Gammaproteobacteria*, especially more OTUs of the orders *Cellvibrionales, Vibrionales, Legionellales, Alteromonadales* were increasingly detected in foam compared to the SML and ULW, whereas the order *Oceanospirilliales* was more depleted in foam (Supplementary Figure 7).

### *Gammaproteobacteria* are typical sea foam colonizers

Using the linear discriminant analysis effect size (LefSe) method we could identify OTUs that were significantly more abundant in foam compared to SML and ULW (Figure 6). The analysis does not refer to the most abundant OTUs in terms of absolute numbers but points out the largest differences between foam and the other two habitats. Members of the *Gammaproteobacteria* were typically abundant active and total foam colonizers (Figure 6). Taxa including *Winogradskyella, Vibrio, Halioglobus* and *Pseudoaltermonas* were particularly abundant in both 16S rRNA and 16S rRNA gene derived foam samples as well as when compared to SML and ULW habitats. *Persicirhabdus* and other *Verrucomicrobiaceae* were typical foam-dwellers with 11% and 7% relative abundance according to their presence in 16S rRNA gene-based samples, but with low activity according to 16S rRNA samples.

Typical SML populating bacteria belonged to taxa, which were phylogenetically related to *Alphaproteobacteria, Gammaproteobacteria* and *Flavobacteria*. Strikingly, abundance of total and active foam specific OTUs often indicated a decreasing gradient from foam to SML to ULW (Supplementary Figure 1). High relative abundances (>5%) of *Planktomarina*, SAR116 and SAR86 were found for 16S rRNA and 16S rRNA gene-based SML samples. SAR11 and Candidatus *Actinomarina* typically occurred in high abundances in the ULW.

### *Trichodesmium* sp.-produced foam – a case study

Due to technical restrictions we could only obtain a single sample from the Timor Sea (Station 8) but found it valuable to analyze because the immediate source (*Trichodesmium* sp.-produced foam) was clear. Among the 16S rRNA gene-based community in foam we found most particle-associated OTUs assigned to *Trichodesmium* (relative abundance=33.4%), *Alteromonas* (26.4%) and *Rhodobium* (5.4%), whereas free-living OTUs were mostly assigned to *Alteromonas* (18.0%) and *Rhodobium* (10.2%) (Supplementary Table 4). Particle-associated OTUs were mainly assigned to *Trichodesmium* (68%) and *Rhodobium* (10.9%) in the SML, and to *Trichodesmium* (23.8%) and *Oscillatoria* (26.7%) in the ULW. Bacteria of the genus *Saprospira* were also detected in foam and SML, mainly in the particle-associated fractions. Most free-living OTUs from SML and ULW were assigned to *Synechococcus* with 15.7% and 21.6% relative abundance, respectively. In all 16S rRNA samples, *Trichodesmium* was also the most abundant among active OTUs in foam and SML, only in the ULW *Oscillatoria* (48.2%) had higher relative abundance compared to *Trichodesmium* (29.1%). The relative abundance of 16S rRNA-based OTUs assigned to *Alteromonas* in foams (particle-associated: 17.8%, free-living: 12.6%) was comparatively enhanced to the SML (particle-associated: 0.2%, free-living: 1.2%) (Supplementary Table 4).

## Discussion

### Foams form ecological niches for selected water bacterial communities and opportunistic *Gammaproteobacteria*

Foams are peculiar but understudied microbial habitats at air-water interfaces, and as soon as they subside, their material becomes part of the SML (Kuznetsova and Lee 2002). As foam has been suggested to represent compressed SML (reviewed by Schilling and Zessner (2011)), we hypothesized that foams contain typical SML bacteria. Our analysis showed that a considerable number of OTUs (>1000) overlap between all three habitats (Figure 3B), supporting that bacteria are regularly dispersed between the biomes by wind-induced mixing processes, small-scale turbulences (Wu, *et al*. 2019) and bubble transport (Robinson, *et al*. 2019). However, also the sampling method, being identical for foam and SML, causes some cross-contamination between habitats. As expected for a compressed form of SML, we found high concentrations of SAS (max. 148,233 µg Teq L^-1^) in foams compared to reported values of 4,989 µg Teq L^-1^ in nearshore SML (Wurl, *et al*. 2011). In line with previous reports, we also found a strong enrichment of cells in foam (Kuznetsova and Lee 2002; Rahlff, *et al*. 2017a; Robinson, *et al*. 2019). Decreasing abundance of small eukaryotic phototrophic cells in the SML during simultaneous increase of those in the respective foam sample supports passive transport from SML to foam, and that SML compression is a mechanism for foam formation (Schilling and Zessner 2011). In our study, typical abundant and free-living groups from the ULW including *Planktomarina*, SAR116 and SAR86 were found in foams at lower relative abundance (Figure 6, Supplementary Figure 11) suggesting that these groups were passively transferred to the sticky SML and further into the foam.

The foam environment is an ecological niche with unique properties, that harbors a distinctive bacterial community when compared to the SML and ULW (Figure 4), which contradicts our initial hypothesis that SML and foam communities would be more similar to each other than foam and ULW. The occurrence of specific foam bacterial communities most likely resembles the particularities of the foam habitat, e.g., its dynamics in formation and disruption, its ephemerality, and high organic matter accumulation. Foam consists of 90% air (Napolitano and Cicerone 1999) and contains channels, referred to as Plateau borders (Schramm and Wassmuth 1994) tending to trap living, motile algae, but draining dead ones (Roveillo, *et al*. 2020). If this mechanism also applies to bacteria, it may explain why most OTUs found in foams were also active (Figure 3A). In addition, foam is heavily enriched in nutrients but also in pollutants (Eisenreich, *et al*. 1978; Pojasek and Zajicek 1978) probably affecting its bacterial community structure. The type of foam under investigation also seems to be important: Foams originating from phytoplankton exudates (Supplementary Table 1), contained high loads of SAS (Figure 2C) and were linked to higher abundances of microbes (Figure 2A&B) compared to foams formed by convergence of surface water (Figure 1C, Supplementary Table 1). However, the role of the foam source for shaping its microbiome requires further research.

The overall difference in the bacterial community composition of foams compared to other habitats (Figure 4) could be due to bacteria originating from SML and ULW that were selectively enriched in foam, including *Vibrio, Pseudoalteromonas*, and *Halioglobus* (Figure 6). Foams associated with phytoplankton biomass likely contain substantial amounts of labile organic matter and nutrients. Labile organic matter particularly selects for fast-growing, opportunistic, and active bacteria in foams such as *Gammaproteobacteria* (Landa, *et al*. 2013; Teeling, *et al*. 2012), which are known to be dominant in the SML (Sun, *et al*. 2020) and on transparent exopolymer particles (Zäncker, *et al*. 2019). For instance, *Alteromonas* sp., of which we found an OTU among the free-living bacteria present in *Trichodesmium* foam (Supplementary Table 4), were previously shown to be highly abundant and active degraders of alginate, a cell wall component from marine macroalgae (Mitulla, *et al*. 2016), and of labile dissolved organic carbon (Pedler, *et al*. 2014).

### The role of particles for foam-populating bacteria and biogenic foam formation

Foams in aquatic habitats contain large numbers of benthic or symbiotic rather than free-living bacteria (Maynard 1968). Since particulate organic matter is frequently enriched in both the SML (Aller, *et al*. 2005) and foams (Johnson, *et al*. 1989) compared to the ULW, foams are ideal substrates for particle-attached colonization. Particle-associated and free-living bacteria form separate communities in many aquatic environments (Crespo, *et al*. 2013; Crump, *et al*. 1999) including all habitats from this study, especially foams (Figure 4). SML bacteria are rather attached to substrates than occurring in the free-living state (Cunliffe, *et al*. 2009), and particle-associated bacteria of the SML are generally more prone to changes in community composition than free-living ones (Stolle, *et al*. 2010). In agreement with that and former studies (Parveen, *et al*. 2013; Rieck, *et al*. 2015), we found higher OTU numbers present on particles independent of the habitat under investigation (Figure 3A).

Although attachment to particles might have some drawbacks for bacteria regarding grazing (Albright, *et al*. 1987), disadvantages might be easily outweighed by benefits of particles providing organic material and shelter from extreme levels of ultraviolet and solar radiation experienced at the air-sea boundary. The LefSe analysis revealed that *Winogradskyella* (*Flavobacteriaceae*), often being associated with brown algae or sponges (Park and Yoon 2013; Schellenberg, *et al*. 2017; Yoon and Lee 2012), was abundant among the metabolically active OTUs in foams (Figure 6). As broken algal cells and detritus are major parts of foams, high relative abundance of *Winogradskyella* in the foam particle-associated fraction (Supplementary Figure 11) might be due to its attachment to algal-derived substrates. *Persicirhabdus*, known for particle adherence in sediments (Freitas, *et al*. 2012) or on plastic debris (Oberbeckmann, *et al*. 2016), was abundant in particle-attached total communities in foam but not very active (Figure 6). Both *Persicirhabdus* and *Winogradskyella* are well-known for their polysaccharide-degrading capacities (Cardman, *et al*. 2014; Yoon and Lee 2012) and, hence, might prefer to stick to organic materials feeding or sheltering them. Microbes with a free-living lifestyle, such as *Planktomarina*, for example, represent a very prominent and active group of the *Rhodobacteraceae* in marine temperate and (sub-)polar areas of the oceans and coastal waters (Giebel, *et al*. 2009; Giebel, *et al*. 2011; Selje, *et al*. 2004; Voget, *et al*. 2015). *Planktomarina* is not a typical biofilm colonizer, but due to its high abundances in surface waters, we assume it was “glued” in the SML and foam fraction and can benefit from enriched dissolved organic material, from aerobic anoxygenic photosynthesis, and oxidation of carbon monoxide as complementary energy sources (Giebel, *et al*. 2019).

Foam events were observed in association with blooms of the haptophyte *Phaeocystis globosa* or *P. pouchettii, Cyanobacteria*, or linked to certain plants (Eberlein, *et al*. 1985; Seuront, *et al*. 2006; Velimirov 1980; Wegner and Hamburger 2002; Wu, *et al*. 2019). Experiments by Velimirov (1980) demonstrated foam formation in the presence of *Ecklonia maxima* while most bacterial growth was antibiotically inhibited. The author showed that an algal component was crucial for the production of stable foam. Nevertheless, foam bacteria might contribute to the foam formation process (Heard, *et al*. 2008), because, like phytoplankton, they produce SAS and exopolysaccharides (Satpute, *et al*. 2010). Foam samples from our study contained bacterial OTUs likely capable of producing SAS, as previously demonstrated for *Vibrio* and *Pseudoalteromonas* (Dang, *et al*. 2016; Hu, *et al*. 2015). Moreover, *Saprospira* found in *Trichodesmium*-associated foam can produce sticky substances such as acidic polysaccharides, which enhance aggregate formation (Furusawa, *et al*. 2015). We further speculate that in the Jade Bay, foam formation could be additionally supported by blooming *Phaeocystis* sp. (OSPAR_Assessments_HASEC17/D503 2017). Surface scum formation by cyanobacteria using intracellular gas vacuole formation as described for *Trichodesmium* sp. (Walsby 1992) could be an ecological strategy to reach atmospheric carbon dioxide supplies at the air-sea interface (Paerl and Ustach 1982). *Trichodesmium* sp. was active in our Timor Sea samples (Supplementary Table 4). During research in the Timor Sea, we incubated surface water with 1 mL *Trichodesmium* foam and found complete oxygen depletion after <14 hours. Since samples without foam showed incomplete O2 consumption (Rahlff, *et al*. 2017b), we assume that complete O2 depletion was attributable to highly active, heterotrophic bacteria that accompanied *Trichodesmium* and their associated foam.

Overall, our cell count and SAS data support the previous assumption that foams are strongly influenced by the SML and, as an ephemeral and discrete habitat, select for specific bacteria including the typical SML inhabitants *Vibrio* and *Pseudoalteromonas*. Moreover, ultramicrobacteria in foams can probably be easily be aerosolized to the atmosphere and dispersed to land (Rahlff, *et al*. 2020). Future studies should elucidate how wind-stirred bulk water and bubbles transfer colonies to SML and foams, and whether bacteria use these habitats as nutrient-rich “rest stop” before being transferred to sea-spray aerosols, dispersed to beaches or returned to bulk water. The SML spans 71% of the Earth’s surface and much remains to be learned about patchy surface phenomena such as foams and their ecological implications for the functioning of the marine food web and carbon turnover.

## Funding

This work was supported by the ERC project PASSME [grant number GA336408] as well as by the Leibniz Association project MarParCloud [grant number SAW-2016-TROPOS-2]. DPRH was supported by the European Regional Development Fund/Estonian Research Council funded Mobilitas Pluss Top Researcher grant “MOBTT24”.

### Acknowledgements

We thank the captain and crew of the *R/V* Falkor during cruise FK161010. We highly appreciate the help of Julius Schmidt and the ICBM workshop team for arranging small boat operations in the Jade Bay, North Sea. We are also grateful to our colleagues Isabel Stahmer, Tiera-Brandy Robinson, Kimberley Bird and Mariana Ribas-Ribas for their assistance during water sampling. Further we appreciate the expert technical assistance of Lisa Maria Engl and Mathias Wolterink.

