## Supplementary Material for "Sea foams are ephemeral hotspots for distinctive bacterial communities contrasting sea-surface microlayer and underlying surface water"

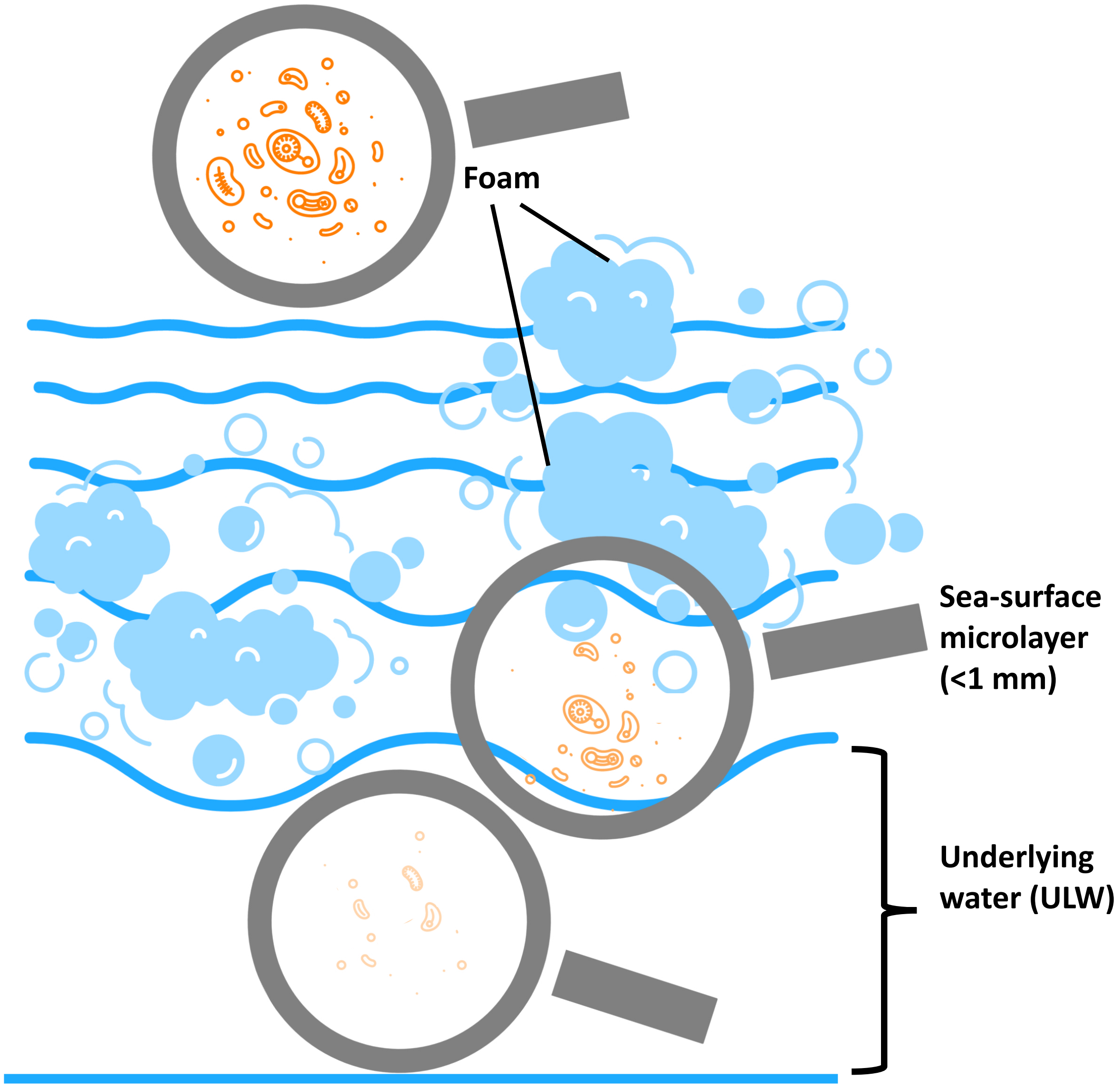


**Supplementary Figure 1:** Concept of the sea-surface microlayer (SML) and floating foams showing decreasing prokaryotic cell abundance from foam over SML to underlying water.

**A**

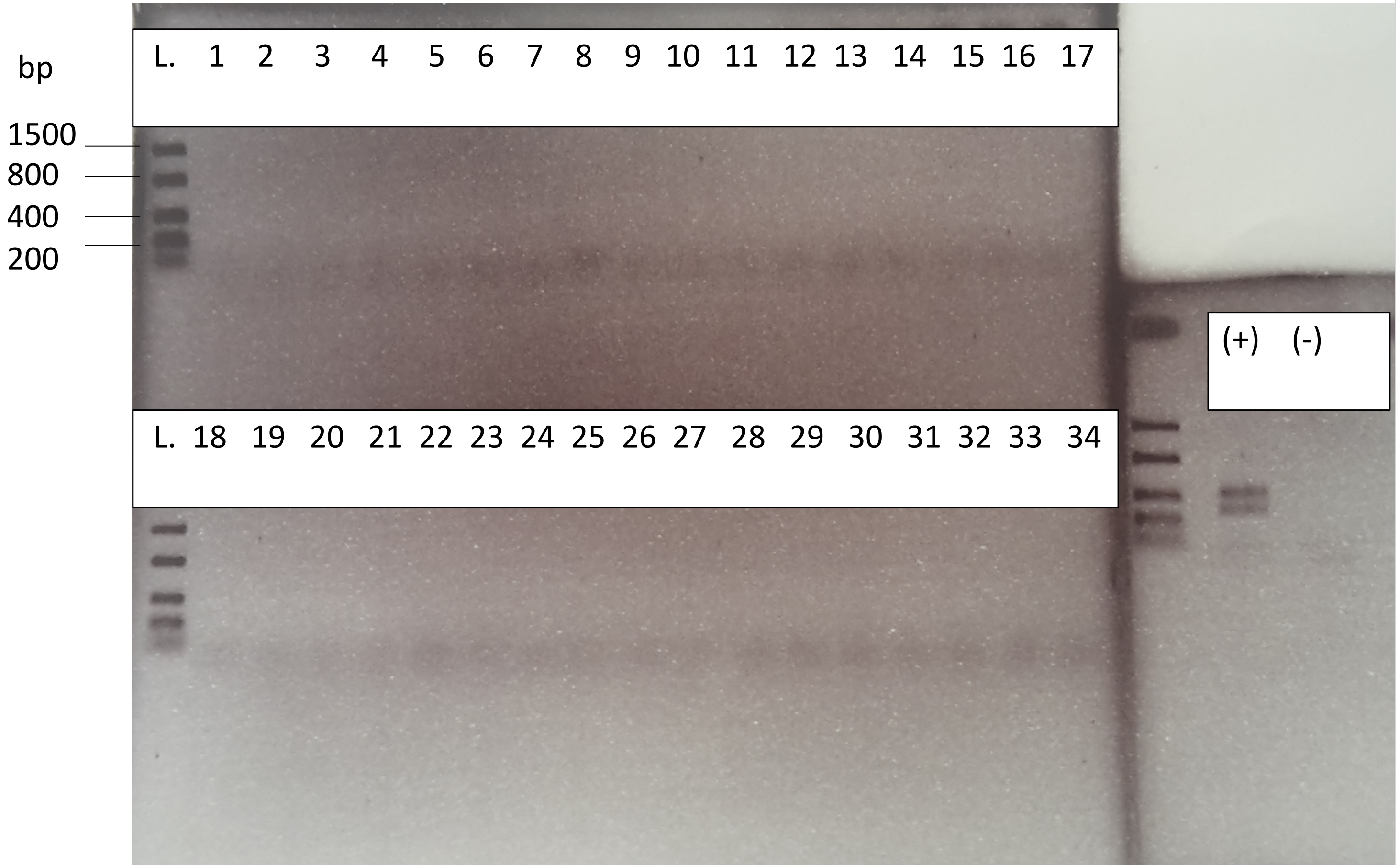


| No | SampleType | SampleName |
|---|---|---|
| 1 | RNA | Fo1_210416_3µ |
| 2 | RNA | SML1_210416_3µ |
| 3 | RNA | ULW1_210416_3µ |
| 4 | RNA | Fo2_210416_3µ |
| 5 | RNA | Fo1_190516_3µ |
| 6 | RNA | SML1_190516_3µ |
| 7 | RNA | ULW1_190516_3µ |
| 8 | RNA | Fo1_190516_0.2µ |
| 9 | RNA | SML1_190516_0.2µ |
| 10 | RNA | ULW1_190516_0.2µ |
| 11 | RNA | Fo3_190516_3µ |
| 12 | RNA | SML3_190516_3µ |
| 13 | RNA | ULW3_190516_3µ |
| 14 | RNA | Fo3_190516_0.2µ |
| 15 | RNA | SML3_190516_0.2µ |
| 16 | RNA | ULW3_190516_0.2µ |
| 17 | RNA | Fo1_190716_3µ |
| 18 | RNA | SML1_190716_3µ |
| 19 | RNA | ULW1_190716_3µ |
| 20 | RNA | Fo1_190716_0.2µ |
| 21 | RNA | SML1_190716_0.2µ |
| 22 | RNA | ULW1_190716_0.2µ |
| 23 | RNA | Fo2_190716_3µ |
| 24 | RNA | SML2_190716_3µ |
| 25 | RNA | ULW2_190716_3µ |
| 26 | RNA | Fo2_190716_0.2µ |
| 27 | RNA | SML2_190716_0.2µ |
| 28 | RNA | ULW2_190716_0.2µ |
| 29 | RNA | Fo_St.8_191016_3µ |
| 30 | RNA | SML_St.8_191016_3µ |
| 31 | RNA | ULW_St.8_191016_3µ |
| 32 | RNA | Fo_St.8_191016_0.2µ |
| 33 | RNA | SML_St.8_191016_0.2µ |
| 34 | RNA | ULW_St.8_191016_0.2µ |

**B**

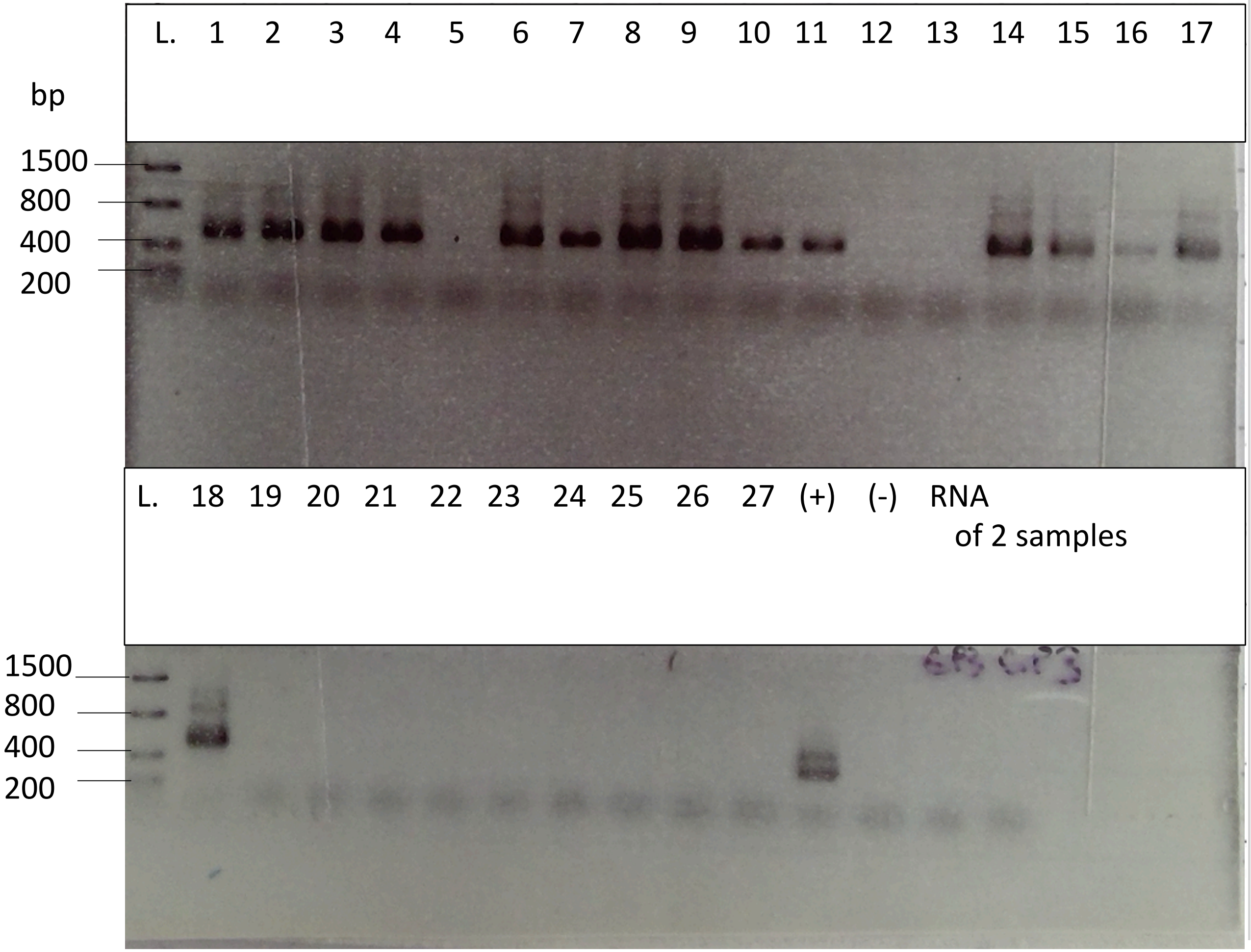


| No | SampleType | SampleName |
|---|---|---|
| 1 | RNA transcribed +RT | Fo1_210416_3µ |
| 2 | RNA transcribed +RT | SML1_210416_3µ |
| 3 | RNA transcribed +RT | ULW1_210416_3µ |
| 4 | RNA transcribed +RT | Fo2_210416_3µ |
| 5 | RNA transcribed +RT | SML2_210416_3µ |
| 6 | RNA transcribed +RT | ULW2_210416_3µ |
| 7 | RNA transcribed +RT | Fo1_190516_3µ |
| 8 | RNA transcribed +RT | SML1_190516_3µ |
| 9 | RNA transcribed +RT | ULW1_190516_3µ |
| 10 | RNA transcribed +RT | Fo1_190516_0.2µ |
| 11 | RNA transcribed +RT | SML1_190516_0.2µ |
| 12 | RNA transcribed +RT | ULW1_190516_0.2µ |
| 13 | RNA transcribed +RT | Fo3_190516_3µ |
| 14 | RNA transcribed +RT | SML3_190516_3µ |
| 15 | RNA transcribed +RT | ULW3_190516_3µ |
| 16 | RNA transcribed +RT | Fo3_190516_0.2µ |
| 17 | RNA transcribed +RT | SML3_190516_0.2µ |
| 18 | RNA transcribed +RT | ULW3_190516_0.2µ |
| 19 | RNA transcribed -RT | Fo1_210416_3µ |
| 20 | RNA transcribed -RT | ULW1_210416_3µ |
| 21 | RNA transcribed -RT | Fo2_210416_3µ |
| 22 | RNA transcribed -RT | ULW2_210416_3µ |
| 23 | RNA transcribed -RT | Fo1_190516_3µ |
| 24 | RNA transcribed -RT | Fo3_190516_3µ |
| 25 | RNA transcribed -RT | ULW1_190516_3µ |
| 26 | no RNA +RT | Control |
| 27 | RNA transcribed -RT | SML2_210416_3µ |

**C**

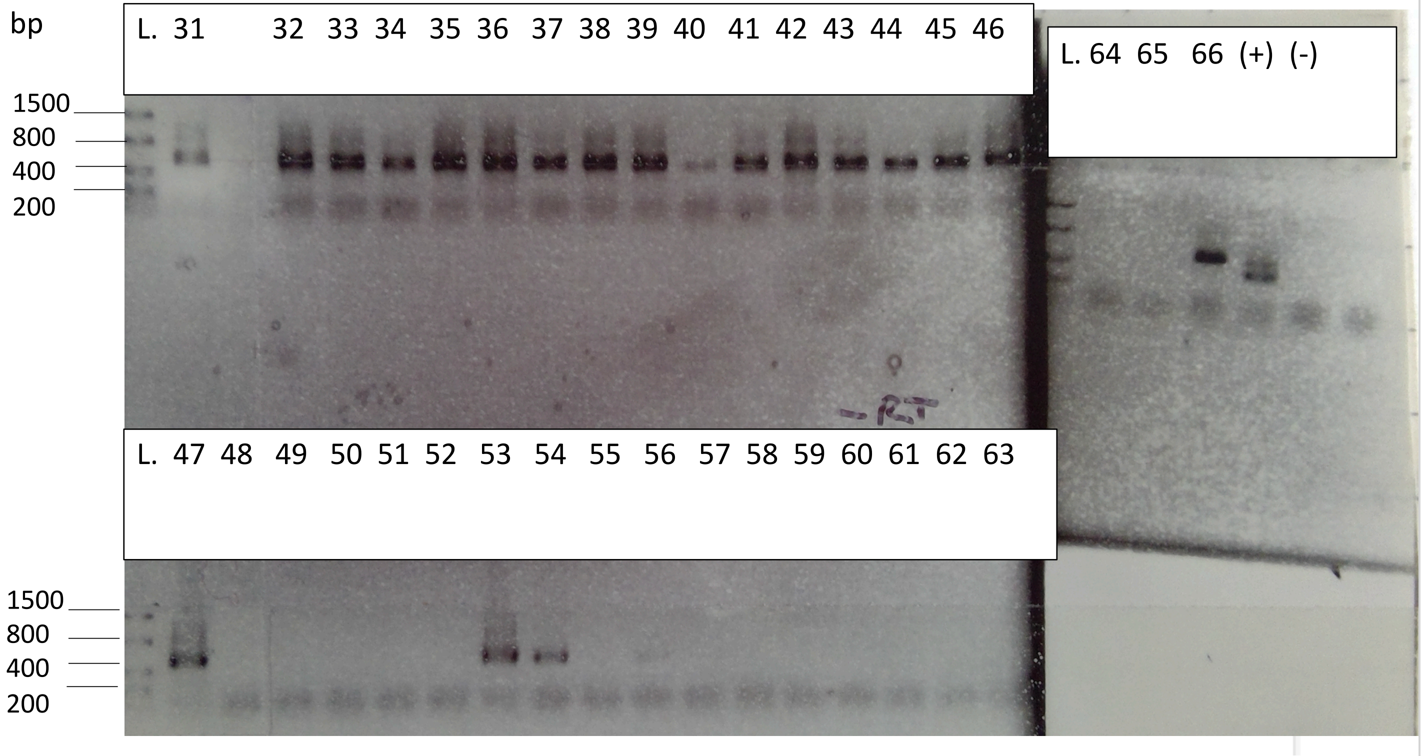


| | | |
|---|---|---|
| 31 | RNA transcribed +RT | Fo1_190716_3µ |
| 32 | RNA transcribed +RT | SML1_190716_3µ |
| 33 | RNA transcribed +RT | ULW1_190716_3µ |
| 34 | RNA transcribed +RT | Fo1_190716_0.2µ |
| 35 | RNA transcribed +RT | SML1_190716_0.2µ |
| 36 | RNA transcribed +RT | ULW1_190716_0.2µ |
| 37 | RNA transcribed +RT | Fo2_190716_3µ |
| 38 | RNA transcribed +RT | SML2_190716_3µ |
| 39 | RNA transcribed +RT | ULW2_190716_3µ |
| 40 | RNA transcribed +RT | Fo2_190716_0.2µ |
| 41 | RNA transcribed +RT | SML2_190716_0.2µ |
| 42 | RNA transcribed +RT | ULW2_190716_0.2µ |
| 43 | RNA transcribed +RT | Fo_St.8_191016_3µ |
| 44 | RNA transcribed +RT | SML_St.8_191016_3µ |
| 45 | RNA transcribed +RT | ULW_St.8_191016_3µ |
| 46 | RNA transcribed +RT | Fo_St.8_191016_0.2µ |
| 47 | RNA transcribed +RT | SML_St.8_191016_0.2µ |
| 48 | RNA transcribed +RT | ULW_St.8_191016_0.2µ |
| 49 | RNA transcribed +RT | repeat sample 5 |
| 50 | RNA transcribed +RT | repeat sample 5 1:10 |
| 51 | RNA transcribed +RT | repeat sample 12 |
| 52 | RNA transcribed +RT | repeat sample 12 1:10 |
| 53 | RNA transcribed +RT | repeat sample 13 |
| 54 | RNA transcribed +RT | repeat sample 13 1:10 |
| 55 | RNA transcribed +RT | repeat sample 13 1:100 |
| 56 | RNA transcribed +RT | repeat sample 16 |
| 57 | RNA transcribed +RT | repeat sample 16 1:10 |
| 58 | RNA transcribed -RT | Fo1_190716_3µ |
| 59 | RNA transcribed -RT | ULW1_190716_3µ |
| 60 | RNA transcribed -RT | Fo2_190716_3µ |
| 61 | RNA transcribed -RT | ULW2_190716_3µ |
| 62 | RNA transcribed -RT | Fo_St.8_191016_3µ |
| 63 | RNA transcribed -RT | Fo_St.8_191016_0.2µ |
| 64 | RNA transcribed -RT | Fo3_190516_3µ |
| 65 | no RNA +RT | Control |
| 66 | RNA transcribed +RT | Fo3_190516_3µ |

**Supplementary Figure 2:** Agarose gel pictures showing A) contamination-free RNA by missing amplicons after polymerase chain reaction (PCR), B) 16S rRNA amplicons after template/cDNA synthesis with reverse transcriptase (+RT) and lack of bands in representative samples after synthesis without (-RT) for PCR samples 1-27, and C) 31-66; Lack of quality of gel pictures is due to picturing of photos from the laboratory journal. (+)=positive control, (-)=negative(no template) control; DNA Ladder (L.) is the FastRuler low range (Thermo Scientific, Cat#SM1103).

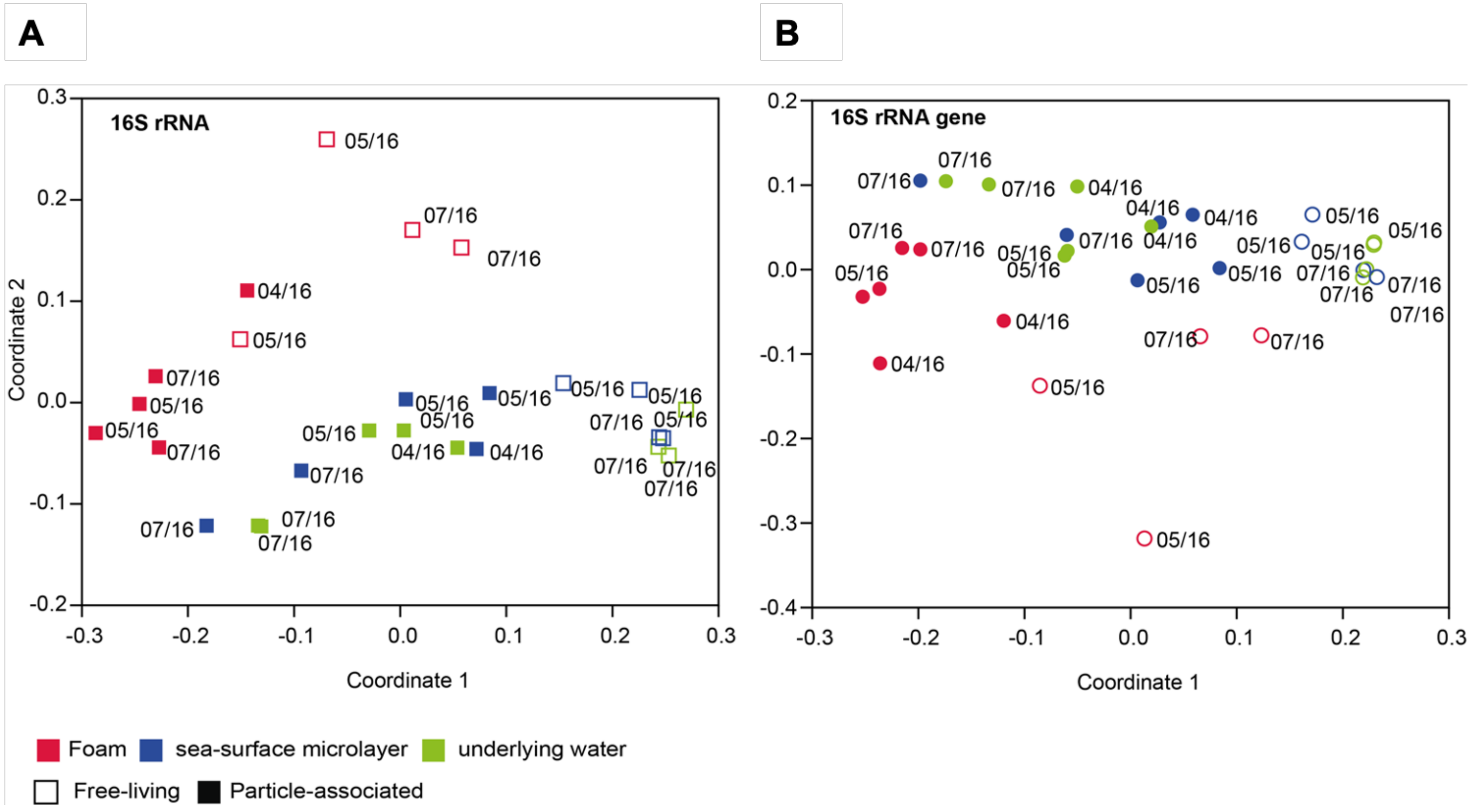


**Supplementary Figure 3:** Non-metric multidimensional scaling plot shows distinct clustering of foam (red), sea-surface microlayer (blue) and underlying water (green) bacterial communities according to sampling dates based on A) 16S rRNA (stress=0.136) and B) 16S rRNA gene-derived amplicons (stress=0.113).

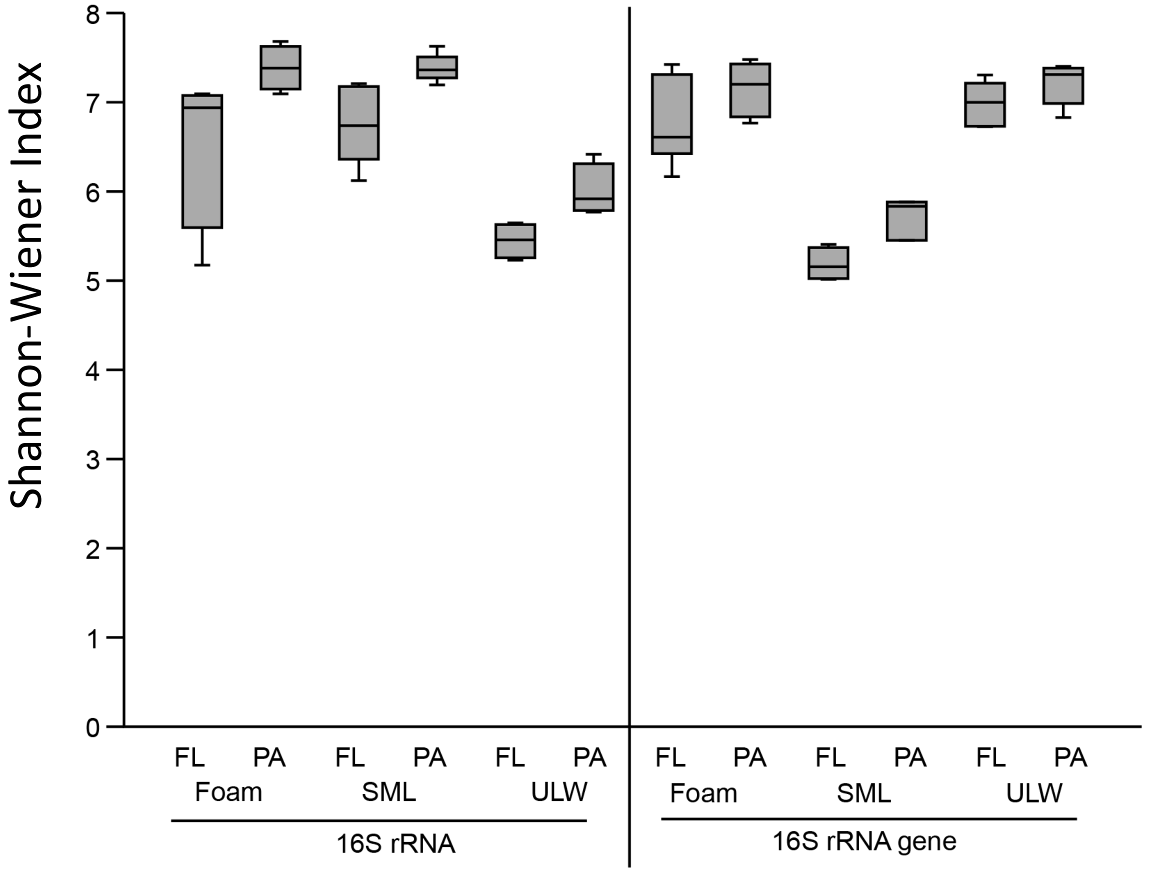


**Supplementary Figure 4:** Shannon-Wiener index reflecting abundance and evenness of North Sea-derived OTUs separated by habitat (foam, sea-surface microlayer (SML) and underlying water (ULW) and by attachment status (free-living (FL) or particle-attached (PA)). The analysis was separated for 16S rRNA and its gene reflecting active and total OTUs, respectively.

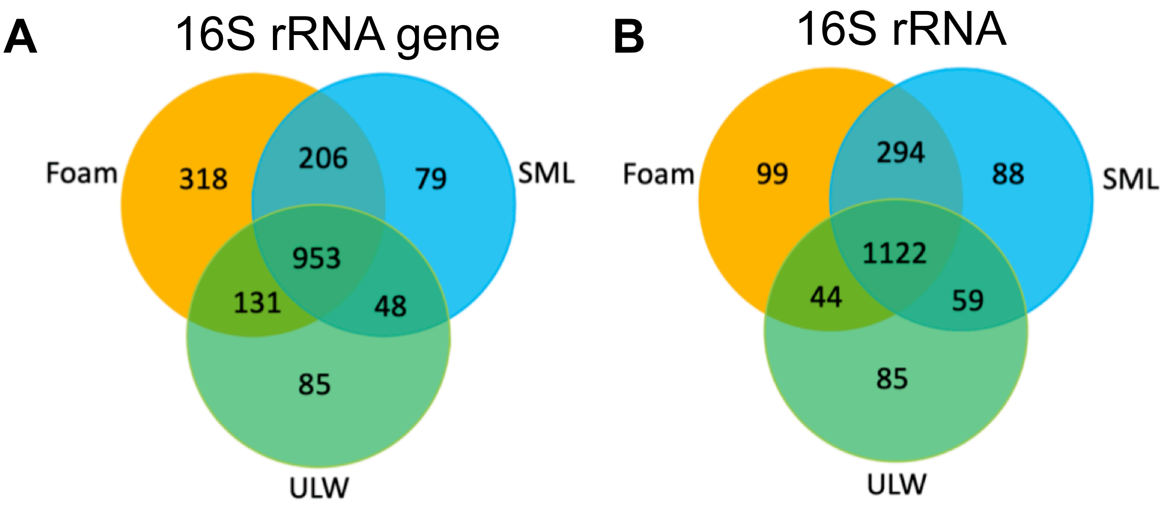


**Supplementary Figure 5:** 3-Venn diagram showing overlapping and unique A) DNA-based and B) cDNA-based OTUs for foam, sea-surface microlayer (SML) and underlying water (ULW). The analysis was separated for 16S rRNA and its gene reflecting active and total OTUs, respectively.

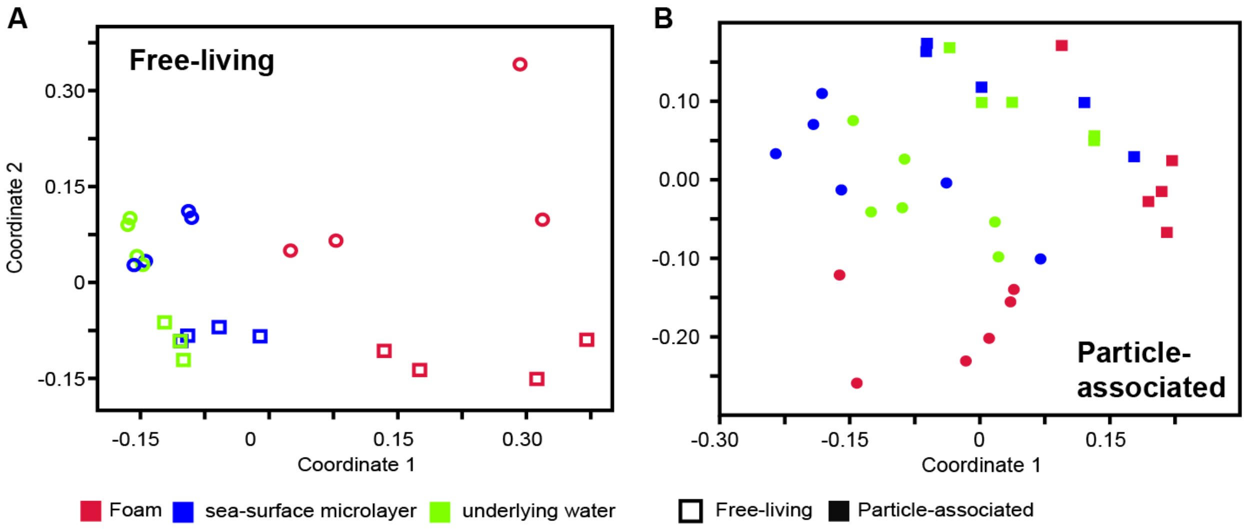


**Supplementary Figure 6:** Non-metric multidimensional scaling plot shows distinct clustering of foam (red), sea-surface microlayer (blue) and underlying water (green) bacterial communities. Further separation of communities into A) free-living with different nucleic acid source (16S rRNA=squares and 16S rRNA gene=circles) stress=0.11; and B) particle-associated with different nucleic acid source (16S rRNA=squares and 16S rRNA gene=circles) stress=0.15.

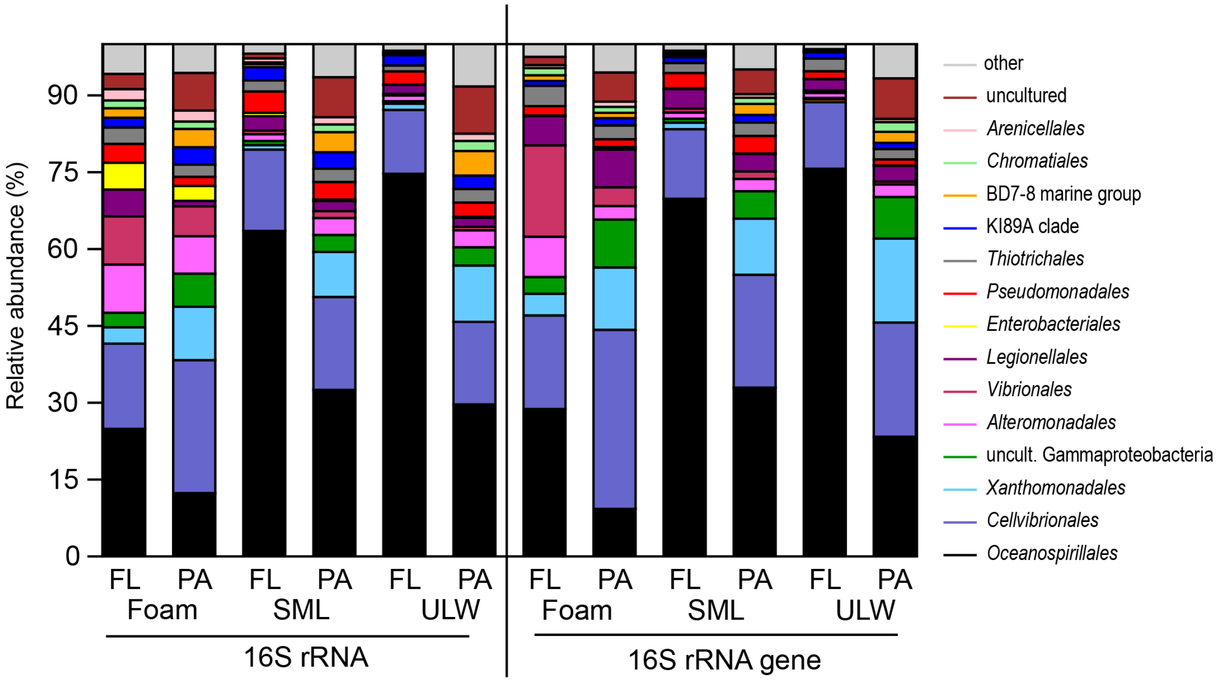


**Supplementary Figure 7:** Beta diversity among *Gammaproteobacteria* in foam, sea-surface microlayer (SML) and underlying water (ULW) samples of 16S rRNA and 16S rRNA gene-based operational taxonomic units (OTUs). Each habitat contains further information on free-living (FL) and particle-associated (PA) bacterial community composition.

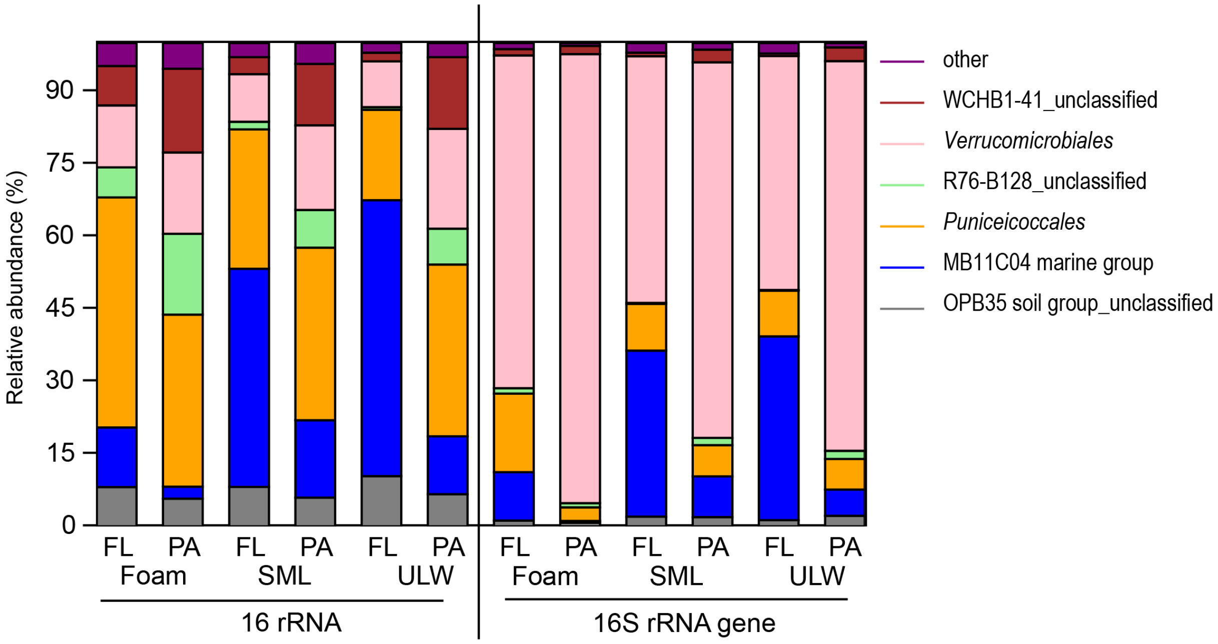


**Supplementary Figure 8:** Beta diversity among *Verrucomicrobia* in foam, sea-surface microlayer (SML) and underlying water (ULW) samples of 16S rRNA and 16S rRNA gene-based operational taxonomic units (OTUs). Each habitat contains further information on free-living (FL) and particle-associated (PA) bacterial community composition.

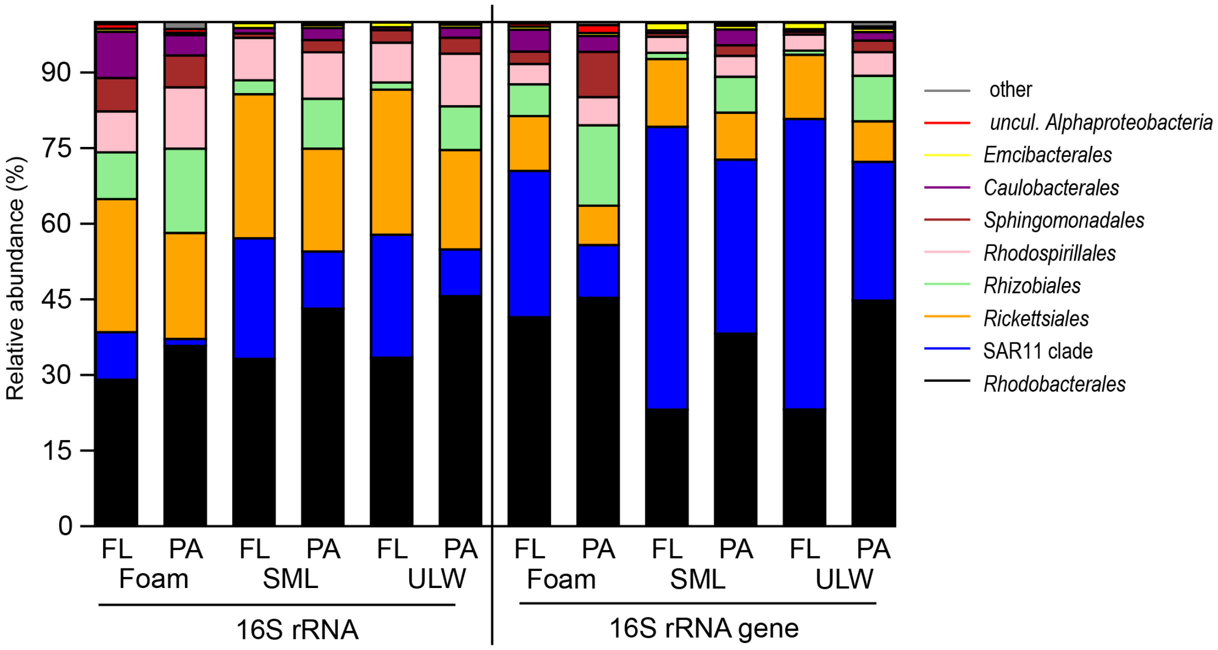


**Supplementary Figure 9:** Beta diversity among *Alphaproteobacteria* in foam, sea-surface microlayer (SML) and underlying water (ULW) samples of 16S rRNA and 16S rRNA gene-based operational taxonomic units (OTUs). Each habitat contains further information on free-living (FL) and particle-associated (PA) bacterial community composition.

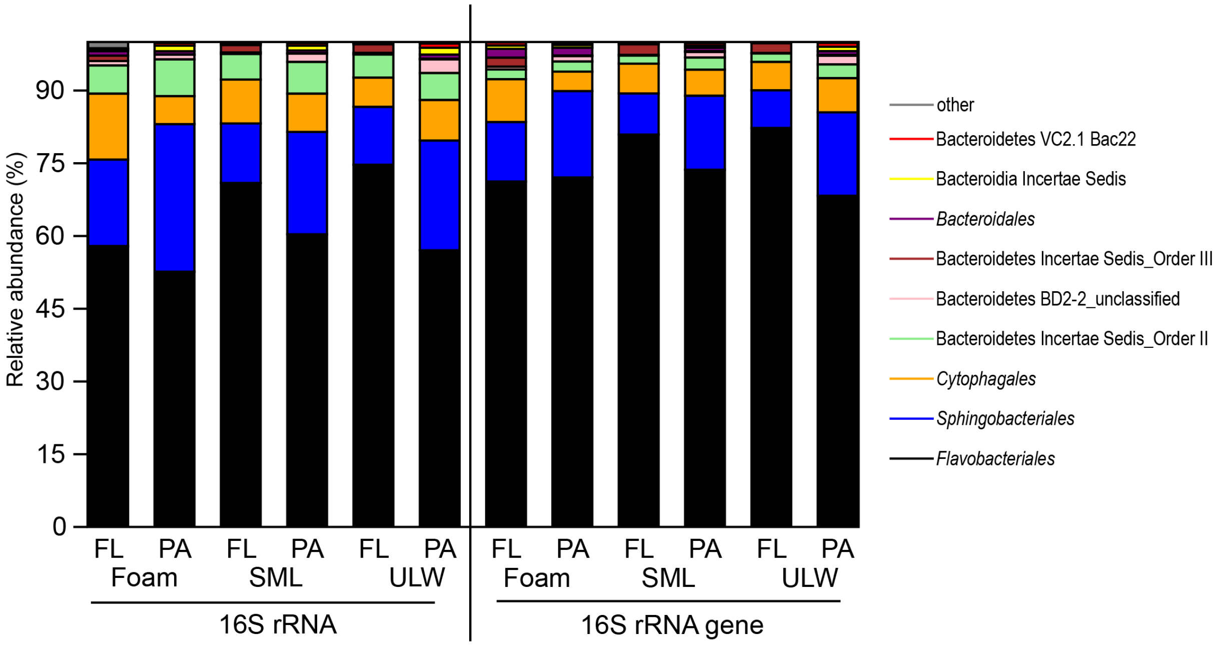


**Supplementary Figure 10:** Beta diversity among *Bacteroidetes* in foam, sea-surface microlayer (SML) and underlying water (ULW) samples of 16S rRNA and 16S rRNA gene-based operational taxonomic units (OTUs). Each habitat contains further information on free-living (FL) and particle-associated (PA) bacterial community composition.

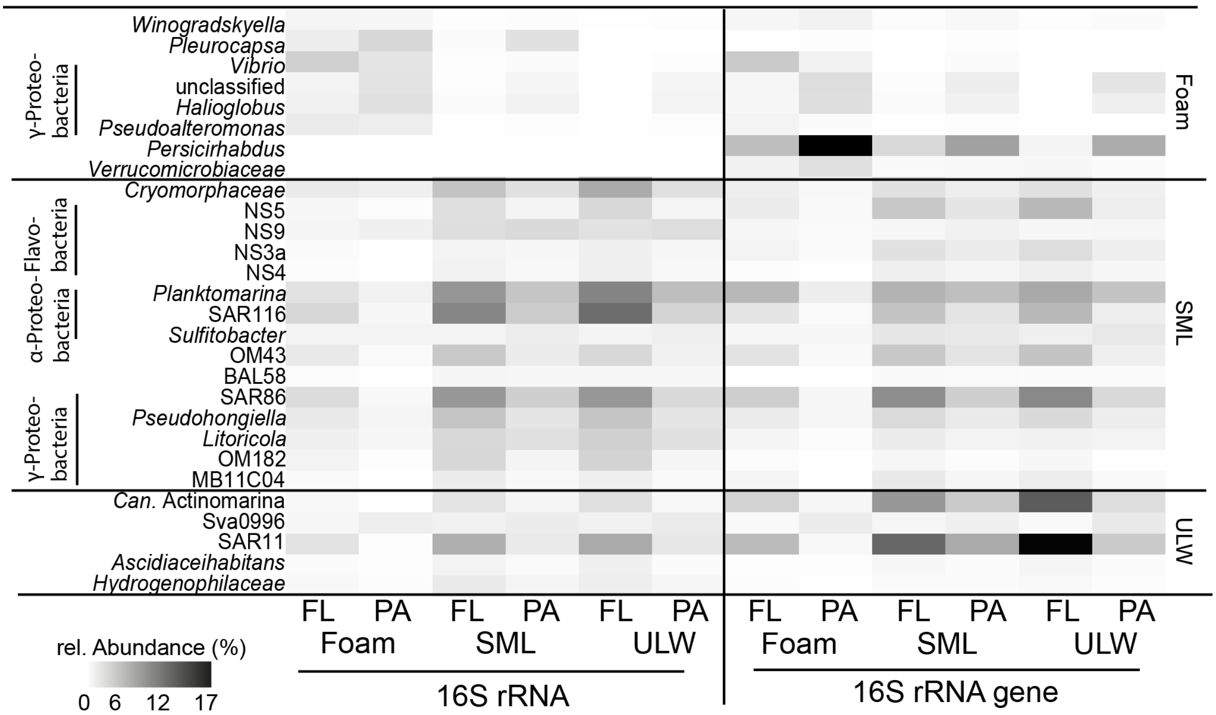


**Supplementary Figure 11:** Heat-map showing the relative abundance of most different foam OTUs compared to sea-surface microlayer (SML) and underlying water (ULW) among free-living (FL) and particle-attached (PA) fractions according to the linear discriminant analysis effect size (LEfSe) method (Segata*, et al.* 2011). The analysis was separated for 16S rRNA and its gene reflecting active and total OTUs, respectively.

**Supplementary Table 1:** Sampling position, sample notes and environmental data; sea-surface microlayer (SML) and underlying water (ULW), n.d.= not determined, NS=North Sea, PAR= Photosynthetically active radiation, rad.=radiation, Temp.=temperature, TS= Timor Sea, UV=ultraviolet

| **Sample set (Foam/SML/ULW)** | **Position** | **Time of sampling (UTC)** | **Remarks** | Wind speed [m s$^{-1}$] | Secchi depth [cm] | Salinity (SML)$^{C}$ | Salinity (ULW)$^{C}$ | Temp. (SML)$^{C}$ | Temp. (ULW)$^{C}$ | Solar rad. [W m$^{-2}$]$^{C}$ | UV index$^{C}$ | PAR [μmol m$^{-2}$ s$^{-1}$]$^{C}$ |
|---|---|---|---|---|---|---|---|---|---|---|---|---|
| NS_FO1_210416 | 53°30.374'N, 08°08.963'E | 13:27 | Foam produced by waves; 0.2 µm filter for nucleic acid extraction had to be discarded. | 4$^{A}$ | 60 | n.d. | n.d. | n.d. | n.d. | 625 | 3.3 | n.d. |
| NS_FO2_210416 | 53°31.089'N, 08°09.998'E | 13:45 | Foam produced by waves; 0.2 µm filter for nucleic acid extraction had to be discarded, no cDNA for this set | 6.7$^{A}$ | 60 | n.d. | n.d. | n.d. | n.d. | 601 | 3.1 | n.d. |
| NS_FO1_190516 | 53°29.916'N, 08°07.9380'E | 09:30 | Slick-associated, probably phytoplankton exudates | <1.0$^{A}$ | 80 | n.d. | n.d. | n.d. | n.d. | 645 | 4.4 | 821 |
| NS_FO3_190516 | n.d. (similar to FO1_190516) | 12:15 | Slick-associated, probably phytoplankton exudates | 1.8$^{A}$ | 80 | n.d. | n.d. | n.d. | n.d. | 307 | 3.5 | 947 |
| NS_FO1_190716 | 53°30.627'N, 08°08.031'E | 10:59 | Close to beach, probably phytoplankton exudates | 1.5$^{A}$ | 60 | 31.7 | 34.7 | 20.1 | 20.0 | n.d. | n.d. | 1662 |
| NS_FO2_190716 | 53°30.327'N, 08°07.854'E | 11:30 | - | 2.5$^{A}$ | 60 | 31.1 | 33.9 | 21.0 | 20.4 | n.d. | n.d. | 1581 |
| TS_FO_St4_151016 | -12°15.49'S, 126°22.36'E | 04:30 | Slick-associated, *Trichodesmium* bloom, only cell counts, no sequencing data | 1.1$^{C}$ | n.d. | 30.0 | 29.8 | 32.2 | 32.0 | 883 | 10.2 | n.d. |
| TS_FO_St5b_171016 | -12°15.46'S, 125°58.60'E | 07:15 | Slick-associated foam, *Trichodesmium* bloom, SML and ULW no slick area, only cell counts, no sequencing data | n.d. | n.d. | n.d. | n.d. | n.d. | n.d. | n.d. | n.d. | n.d. |

| | | | | | | | | | | | | |
|---|---|---|---|---|---|---|---|---|---|---|---|---|
| TS_FO_St8_191016 | -13°41.51'S, 127°31.27'E | 01:05 | *Trichodesmium* bloom, sudden rain and squalls during SML and ULW sampling; No ULW-FL cDNA sample | 4.3[C] | n.d. | 31.6 | 31.1 | 30.3 | 30.5 | 736 | 7.8 | n.d. |

[A] measured with anemometer
[C] measured with catamaran "*Sea Surface Scanner"* and associated sensors (Ribas-Ribas*, et al.* 2017)

**Supplementary Table 2:** List of all pairwise comparisons: Z-statistic (p-value) for Figure 3A based on Dunn's test (Dinno and Dinno 2017) in R version 4.0.3 (Team 2017). Asterisks indicate the level of significant differences: * p≤0.05, ** p≤0.01, *** p≤0.001; Yellow marked comparisons are the ones indicated in Figure 3A because they refer to intra-habitat or intra-attachment status comparisons. Blue comparisons are also significant but compare between different habitats and attachment status and are not shown in Figure 3A. SML=sea-surface microlayer, ULW=underlying water, PA=particle-attached, FL=free-living

| 16S rRNA | Significance | 16S rRNA gene | Significance |
|---|---|---|---|
| foam_FL - foam_PA : -0.877207 (0.3804) | | foam_FL - foam_PA : -2.610318 (0.0090) | ** |
| foam_FL - SML_FL : 1.941782 (0.0522) | | foam_FL - SML_FL : 0.522092 (0.6016) | |
| foam_PA - SML_FL : 2.924026 (0.0035) | ** | foam_PA - SML_FL : 3.182242 (0.0015) | ** |
| foam_FL - SML_PA : -0.389870 (0.6966) | | foam_FL - SML_PA : -0.733235 (0.4634) | |
| foam_PA - SML_PA : 0.516899 (0.6052) | | foam_PA - SML_PA : 2.098642 (0.0358) | * |
| SML_FL - SML_PA : -2.436688 (0.0148) | * | SML_FL - SML_PA : -1.305159 (0.1918) | |
| foam_FL - ULW_FL : 2.140168 (0.0323) | * | foam_FL - ULW_FL : 0.763058 (0.4454) | |
| foam_PA - ULW_FL : 3.044008 (0.0023) | ** | foam_PA - ULW_FL : 3.446207 (0.0006) | *** |
| SML_FL - ULW_FL : 0.342426 (0.7320) | | SML_FL - ULW_FL : 0.240965 (0.8096) | |
| SML_PA - ULW_FL : 2.596360 (0.0094) | ** | SML_PA - ULW_FL : 1.569124 (0.1166) | |
| foam_FL - ULW_PA : 0.194935 (0.8454) | | foam_FL - ULW_PA : -1.114518 (0.2651) | |
| foam_PA - ULW_PA : 1.137179 (0.2555) | | foam_PA - ULW_PA : 1.672355 (0.0945) | |
| SML_FL - ULW_PA : -1.851883 (0.0640) | | SML_FL - ULW_PA : -1.686441 (0.0917) | |
| SML_PA - ULW_PA : 0.620279 (0.5351) | | SML_PA - ULW_PA : -0.426286 (0.6699) | |
| ULW_FL - ULW_PA : -2.059182 (0.0395) | * | ULW_FL - ULW_PA : -1.950406 (0.0511) | |

**Supplementary Table 3:** Relative abundance (%) of operational taxonomic units as shown in Figure 5. SML=sea-surface microlayer, ULW=underlying water, PA=particle-attached, FL=free-living

| **16S rRNA gene** | **Foam_PA** | **Foam_FL** | **SML_PA** | **SML_FL** | **ULW_PA** | **ULW_FL** |
|---|---|---|---|---|---|---|
| >*Gammaproteobacteria* | 25.98 | 22.70 | 22.85 | 18.91 | 24.33 | 17.05 |
| >*Alphaproteobacteria* | 12.31 | 27.65 | 24.98 | 39.18 | 20.56 | 41.33 |
| *Bacteroidetes* | 14.26 | 12.77 | 16.21 | 15.85 | 13.93 | 15.71 |
| *Verrucomicrobia* | 24.86 | 9.09 | 10.23 | 5.26 | 9.99 | 3.28 |
| *Actinobacteria* | 4.51 | 12.70 | 7.36 | 9.92 | 8.92 | 13.01 |
| >*Deltaproteobacteria* | 5.80 | 3.76 | 5.36 | 1.22 | 7.63 | 1.06 |
| >*Betaproteobacteria* | 1.04 | 2.82 | 2.90 | 5.30 | 2.03 | 4.93 |
| *Planctomycetes* | 3.96 | 1.54 | 3.16 | 0.64 | 4.87 | 0.42 |
| *Cyanobacteria* | 0.52 | 1.10 | 0.67 | 0.33 | 0.55 | 0.25 |
| *Gemmatimonadetes* | 0.48 | 0.27 | 0.47 | 0.11 | 0.60 | 0.08 |
| other | 5.50 | 5.22 | 5.40 | 3.00 | 6.11 | 2.57 |
| unclassified | 0.80 | 0.39 | 0.40 | 0.29 | 0.48 | 0.32 |

| **16S rRNA** | **Foam_PA** | **Foam_FL** | **SML_PA** | **SML_FL** | **ULW_PA** | **ULW_FL** |
|---|---|---|---|---|---|---|
| >*Gammaproteobacteria* | 34.98 | 37.40 | 28.59 | 27.83 | 30.14 | 24.94 |
| >*Alphaproteobacteria* | 16.56 | 25.00 | 23.33 | 33.91 | 22.31 | 38.66 |
| *Bacteroidetes* | 18.38 | 13.64 | 16.28 | 16.93 | 16.52 | 19.08 |
| *Verrucomicrobia* | 3.08 | 3.91 | 3.48 | 2.85 | 3.81 | 1.98 |
| *Actinobacteria* | 1.66 | 1.94 | 2.88 | 3.67 | 2.77 | 3.70 |
| >*Deltaproteobacteria* | 7.95 | 3.64 | 7.08 | 1.76 | 8.95 | 1.36 |
| >*Betaproteobacteria* | 1.48 | 3.58 | 3.14 | 6.18 | 2.61 | 4.75 |
| *Planctomycetes* | 2.27 | 0.93 | 1.44 | 0.45 | 1.93 | 0.28 |
| *Cyanobacteria* | 4.79 | 3.02 | 4.55 | 2.31 | 1.13 | 1.67 |
| *Gemmatimonadetes* | 2.31 | 1.35 | 2.28 | 1.13 | 2.21 | 0.78 |
| other | 6.13 | 5.36 | 6.38 | 2.74 | 7.05 | 2.32 |
| unclassified | 0.41 | 0.23 | 0.56 | 0.24 | 0.57 | 0.49 |

**Supplementary Table 4:** Relative abundance (%) of most abundant operational taxonomic units in foam, SML and ULW among free-living (FL) and particle-attached (PA) fractions from Station 8, Timor Sea, separated by 16S rRNA and 16S rRNA gene analysis. The ULW_FL sample for 16S rRNA is missing.

| **16S rRNA gene** | **Foam_PA** | **Foam_FL** | **SML_PA** | **SML_FL** | **ULW_PA** | **ULW_FL** |
|---|---|---|---|---|---|---|
| *Cyanobacteria; Cyanobacteria;* Subsection III; Family I*; Trichodesmium* | 33.39 | 2.11 | 67.96 | 6.17 | 23.78 | 0.02 |
| *Cyanobacteria; Cyanobacteria;* Subsection I; Family I*; Synechococcus* | 3.36 | 8.71 | 0.99 | 15.69 | 4.19 | 21.63 |
| *Proteobacteria; Gammaproteobacteria; Alteromonadales; Alteromonadaceae; Alteromonas* | 26.40 | 18.02 | 1.83 | 2.30 | 3.65 | 2.06 |
| *Proteobacteria; Alphaproteobacteria; Rhizobiales; Rhodobiaceae; Rhodobium* | 5.43 | 10.23 | 10.97 | 2.45 | 8.93 | 0.80 |
| *Cyanobacteria; Cyanobacteria;* Subsection III*;* Family I*; Oscillatoria* | 0.47 | 0.04 | 0.46 | 0.06 | 26.57 | 0.00 |
| *Proteobacteria; Alphaproteobacteria;* SAR11 clade*;* Surface 1 | 0.26 | 1.65 | 0.03 | 7.57 | 0.29 | 8.58 |
| *Cyanobacteria; Cyanobacteria;* Subsection I; Family I*; Prochlorococcus* | 0.30 | 0.93 | 0.06 | 8.10 | 0.48 | 7.50 |
| *Bacteroidetes; Sphingobacteriia; Sphingobacteriales; Saprospiraceae; Saprospira* | 4.52 | 3.36 | 5.90 | 0.02 | 0.99 | 0.01 |
| *Proteobacteria; Gammaproteobacteria; Oceanospirillales;* SAR86 clade | 0.29 | 1.27 | 0.03 | 6.05 | 0.15 | 6.05 |
| *Proteobacteria; Alphaproteobacteria; Rickettsiales;* SAR116 clade | 0.22 | 1.19 | 0.06 | 5.11 | 0.64 | 6.59 |

| **16S rRNA** | **Foam_PA** | **Foam_FL** | **SML_PA** | **SML_FL** | **ULW_PA** |
|---|---|---|---|---|---|
| *Cyanobacteria; Cyanobacteria;* Subsection III; Family I*; Trichodesmium* | 47.44 | 21.71 | 85.44 | 38.84 | 29.08 |
| *Cyanobacteria; Cyanobacteria;* Subsection III; Family I*; Oscillatoria* | 0.65 | 0.49 | 0.06 | 0.36 | 48.15 |
| *Proteobacteria; Alphaproteobacteria; Rhizobiales; Rhodobiaceae; Rhodobium* | 7.96 | 14.44 | 9.17 | 7.20 | 5.77 |
| *Cyanobacteria; Cyanobacteria;* SubsectionI; Family I*; Synechococcus* | 8.33 | 6.44 | 1.26 | 16.05 | 1.92 |
| *Proteobacteria; Gammaproteobacteria; Alteromonadales; Alteromonadaceae; Alteromonas* | 17.66 | 12.63 | 0.16 | 1.22 | 0.57 |
| *Bacteroidetes; Sphingobacteriia; Sphingobacteriales; Saprospiraceae; Saprospira* | 2.88 | 6.63 | 2.00 | 1.37 | 0.90 |
| *Cyanobacteria; Cyanobacteria;* SubsectionI*;* Family I*; Prochlorococcus* | 0.14 | 0.85 | 0.00 | 8.45 | 0.05 |
| *Proteobacteria; Alphaproteobacteria; Rickettsiales;* SM2D12 | 0.67 | 1.98 | 0.03 | 0.93 | 1.70 |
| *Proteobacteria; Alphaproteobacteria; Rhodobacterales; Rhodobacteraceae;*uncultured*;* | 0.22 | 0.40 | 0.02 | 3.08 | 0.42 |
| *Bacteroidetes; Bacteroidetes Incertae Sedis;* Order III*;* Unknown Family*; Balneola;* | 0.56 | 2.04 | 0.35 | 0.50 | 0.42 |
